# Surface growth of *Pseudomonas aeruginosa* reveals a regulatory effect of 3-oxo-C_12_-homoserine lactone in absence of its cognate receptor, LasR

**DOI:** 10.1101/2023.04.14.536888

**Authors:** Thays de Oliveira Pereira, Marie-Christine Groleau, Eric Déziel

## Abstract

Successful colonization of a multitude of ecological niches by the bacterium *Pseudomonas aeruginosa* relies on its ability to respond to concentrations of self-produced signal molecules. This intercellular communication system known as quorum sensing (QS) tightly regulates the expression of virulence determinants and a diversity of survival functions, including those required for social behaviours. In planktonic cultures of *P. aeruginosa*, the transcriptional regulator LasR is generally considered on top of the QS circuitry hierarchy; its activation relies on binding to 3-oxo-C_12_-homoserine lactone (3-oxo-C_12_-HSL), a product of the LasI synthase. Transcription of *lasI* is activated by LasR, resulting in a positive feedback loop. Few studies have looked at the function of QS during surface growth even though *P. aeruginosa* typically lives in biofilm-like communities under natural conditions. Here, we show that surface-grown *P. aeruginosa* readily produces 3-oxo-C_12_-HSL in absence of LasR, and that this phenotype is frequent upon surface association in naturally occurring environmental and clinical LasR-defective isolates, suggesting a conserved alternative function for the signal. Indeed, even in the absence of the cognate regulator LasR, 3-oxo-C_12_-HSL upregulates the autologous expression of pyocyanin and of LasR-controlled virulence determinants in neighboring cells. This highlights a possible role for 3-oxo-C_12_-HSL in shaping community responses and provides a possible evolutive benefit for mixed populations to carry LasR-defective cells, a common feature of natural of *P. aeruginosa*.

**IMPORTANCE:** The bacterium *Pseudomonas aeruginosa* colonizes and thrives in many environments, in which it is typically found in surface-associated polymicrobial communities known as biofilms. Adaptation to this social behavior is aided by quorum sensing (QS), an intercellular communication system pivotal in the expression of social traits. Regardless of its importance in QS regulation, the loss of function of the master regulator LasR is now considered a conserved adaptation of *P. aeruginosa*, irrespective of the origin of strains. By investigating the QS circuitry in surface-grown cells, we found accumulation of QS signal 3-oxo-C_12_-HSL in absence of its cognate receptor and activator, LasR. The current understanding of the QS circuit, mostly based on planktonic growing cells, is challenged by investigating the QS circuitry of surface-grown cells. This provides a new perspective on the beneficial aspects that underline the frequency of LasR-deficient isolates.

## INTRODUCTION

Bacteria are social organisms that often respond to environmental cues in coordination. *Pseudomonas aeruginosa* is a very adaptable Gram-negative bacterium that colonizes diverse ecological niches. The flexibility of this opportunist human pathogen is aided by several regulatory networks, assuring proper responses to changing environmental conditions. Quorum sensing (QS) is a gene expression regulation mechanism based on the production, release, detection and response to diffusible signaling molecules that synchronizes the transcription of target genes in a population density-dependent manner (1). In *P. aeruginosa*, three interlinked QS systems regulate the expression of hundreds of genes – including several encoding virulence determinants (2). In this bacterium, QS regulation is structured as a hierarchical network composed of two *N*-acyl homoserine lactone (AHL)-based circuits (*las* and *rhl*) and the *pqs* system, that relies on signaling molecules of the 4-hydroxy-2-alkylquinoline (HAQ) family. The *las* and *rhl* systems comprise an AHL synthase (LasI and RhlI) responsible for the syntheses of *N*-(3-oxododecanoyl)-L-homoserine lactone (3-oxo-C_12_-HSL) and *N*-butanoyl-L-homoserine lactone (C_4_-HSL), respectively (3, 4). These autoinducers activate their cognate LuxR-type transcriptional regulators – LasR and RhlR, which in turn can induce the transcription of target QS-regulated genes. Under standard laboratory conditions, the *las* system is generally considered to be atop the regulatory hierarchy. Once activated by the binding with its cognate autoinducer, LasR regulates several virulence traits such as the elastase LasB (*lasB*) (5, 6). LasR also induces the transcription of the LasI synthase coding gene, creating a positive feedback loop (7). The *pqs* system relies on the LysR-type transcriptional regulator MvfR (also known as PqsR) (8, 9). The latter directly activates the operons *pqsABCDE* and *phnAB*, both required for HAQ biosynthesis, and indirectly regulates the expression of many other QS-regulated genes via PqsE (8, 10-14). MvfR has dual ligands as it can be induced by 4-hydroxy-2-heptylquinoline (HHQ) and the *Pseudomonas* quinolone signal (PQS; 3,4-dihydroxy-2-alkylquinoline), both members of the HAQ family (15, 16). The *rhl* and *pqs* circuits are directly and positively regulated by LasR, which induces the transcription of *rhlR* and *rhlI* as well as *mvfR* (13, 15, 17, 18).

In addition to sensing the surrounding chemical environment, bacteria are also responsive to mechanical signals, such as those involved in the physical encounter of the cell with surfaces or with each other. Indeed, several behaviors are specific to life on surfaces, including movement on semi-solid (swarming motility) and solid surfaces (twitching motility) as well as biofilm formation (19-21). Not surprisingly, virulence is also induced by surface attachment as many infection strategies require contact with the host (22-24). Even though QS and surface-sensing regulate many of the same social behaviors, little is known about how these different regulatory cues converge to modulate bacterial responses. Exploring the link between surface-sensing and QS is particularly relevant as *P. aeruginosa* readily adopts a surface-attached mode of growth as biofilms in its natural habitats. Biofilms are organized communities encased in a self-produced exopolymeric matrix. In the context of infections, biofilms contribute to host immune evasion and delay antibiotic penetration (25, 26). In fact, *P. aeruginosa* persists as biofilms in the lungs of people with the genetic disease cystic fibrosis (27).

While the emergence of LasR-defective mutants has long been associated with adaptation to the CF lung environment (28-31), it is actually a common feature of *P. aeruginosa* from diverse environments (32, 33). Interestingly, some LasR-defective isolates, known as RAIL (for <u>R</u>hlR <u>a</u>ctive independently of <u>L</u>asR), retain a functional RhlR regulator (31, 32, 34-37). Their sustained QS responses are in line with our previous report showing that in the presence of a nonfunctional LasR, RhlR acts as a surrogate activator for a set of LasR-regulated genes (38). It is noteworthy that in the wild-type *P. aeruginosa* strain PA14 background, surface-sensing upregulates *lasR*, and that surface-grown cells induce LasR targets more strongly than their planktonic counterpart (39). Thus, surface-sensing appears to sensitize cells to the cognate autoinducer 3-oxo-C_12_-HSL. Considering the prevalence of LasR-defective mutants, unable to produce nor respond to 3-oxo-C_12_-HSL, we wondered how *P. aeruginosa* would respond to surface attachment, as biofilm formation is essential to this bacterium physiology and pathology.

In this study, we investigated the effect of surface-sensing on QS responses of LasR-defective strains. We found that, upon surface attachment, LasR becomes dispensable to the production of 3-oxo-C_12_-HSL. This response is conserved among naturally occurring environmental and clinical LasR-defective isolates. Production of 3-oxo-C_12_-HSL modulates the production of virulence factors at the individual (LasR-defective background) and community levels (mixed with LasR-responsive cells). As a result, virulence of mixed populations, composed of LasR-responsive and LasR-defective cells, is accentuated. We propose that the production of 3-oxo-C_12_-HSL by LasR-negative cells, modulating biological bacterial responses on diverse levels, has a positive role in shaping community responses of the population.

## MATERIALS AND METHODS

### Bacterial strains and growth conditions

Bacterial strains and plasmids used in this study are listed in **Table 1** and **Table 2**, respectively. Oligonucleotides used are listed in **Table S1**. Bacteria were routinely grown in tryptic soy broth (TSB; BD Difco, Canada) at 37°C in a TC-7 roller drum (NB, Canada) at 240 rpm or on Lysogeny Broth (LB; BD Difco, Canada) agar plates. For quantification of QS signaling molecules and related data, King’s A broth (planktonic growth) or King’s A agar (surface-associated growth) supplemented with 100 µM FeCl_3_ were used (40). For the latter, sterile King’s A agar was poured into each well of a 96-well plate (200 µl per well) and allowed to solidify at the center of a biosafety cabinet. When needed, the following concentrations of antibiotics were included: for *Escherichia coli* 100 µg/ml carbenicillin, 15 µg/ml gentamicin, and 15 µg/ml tetracycline. Diaminopimelic acid (DAP) was added to cultures of the auxotroph *E. coli* χ7213 at 62.5 μg/ml. Irgasan (20 µg/ml) was used as a counter-selection agent against *E. coli*. For *P. aeruginosa*, 300 µg/ml carbenicillin, 30 µg/ml gentamicin, and tetracycline at 125 µg/ml (solid) or 75 µg/ml (liquid).

**Table 1.** Strains used in this study

| Strain | Lab ID # | Relevant genotype or description | Reference |
| --- | --- | --- | --- |
| *P. aeruginosa* |  |  |  |
| PA14 | ED14 | Clinical isolate from a human burn patient UCBPP-PA14 | (41) |
| PA14 $\Delta lasR$ | ED4409 | PA14 derivative; unmarked in-frame *lasR* deletion | This study |
| PA14 $\Delta lasI$ | ED4539 | PA14 derivative; unmarked in-frame *lasI* deletion | (42) |
| PA14 $\Delta rhlR$ | ED4406 | PA14 derivative; unmarked in-frame *rhlR* deletion | This study |
| PA14 *lasR<sup>-</sup> rhlR<sup>-</sup>* | ED266 | PA14 derivative; marked deletion of *lasR* (*lasR::Gm*) and *rhlR* (*rhlR::Tc*) | (38) |
| PA14 $\Delta lasR \Delta rhlI$ | ED4541 | PA14 derivative; unmarked in-frame double *lasR* and *rhlI* deletion | This study |
| PA14 *lasR<sup>-</sup> $\Delta pqsE$* | ED247 | PA14 derivative; marked deletion of *lasR* (*lasR::Gm*) and an unmarked *pqsE* deletion | (13) |
| PA14 $\Delta lasR \Delta lasI$ | ED4540 | PA14 derivative; unmarked in-frame double *lasR* and *lasI* deletion | This study |
| PA14 $\Delta lasR \Delta lasI \Delta rhlI$ attB::CTX *phzA1-lux* | ED4544 | PA14 derivative; unmarked in-frame triple *lasR*, *lasI* and *rhlI* deletion carrying the chromosomal *phzA1-lux* reporter | This study |
| PA14 $\Delta lasR \Delta lasI$ attB::CTX *phzA1-lux* | ED4591 | PA14 derivative; ED4540 carrying the chromosomal *phzA1-lux* reporter | This study |
| PA14 $\Delta lasI$ attB::CTX *lasB-lux* | ED4543 | PA14 derivative; ED4539 carrying the chromosomal *lasB-lux* reporter | This study |
| PA14 $\Delta lasR$ attB::CTX *lasI-lux* | ED4542 | PA14 derivative; ED4409 carrying the chromosomal *lasI-lux* reporter | This study |
| PA14 $\Delta lasR \Delta pilT$ | ED4556 | PA14 derivative; unmarked in-frame double *lasR* and *pilT* deletion | This study |
| PA14 $\Delta lasR pilU$ | ED4557 | PA14 derivative; unmarked in-frame *lasR* and marked *pilU* mutant (*pilU::MrT7*) | This study |
| 18G | ED4592 | Oil-contaminated soil | (43) |
| 32R | ED4593 | Oil-contaminated soil | (43) |
| 78RV | ED4590 | Oil-contaminated soil | (43) |
| E41 | ED4160 | Cystic fibrosis isolate | (31) |
| E113 | ED4144 | Cystic fibrosis isolate | (31) |
| E167 | ED4152 | Cystic fibrosis isolate | (31) |
| E113 $\Delta rhlR$ | ED4145 | E113 derivative carrying an unmarked deletion in the *rhlR* gene | (37) |
| E167 $\Delta rhlR$ | ED4153 | E167 derivative carrying an unmarked deletion in the *rhlR* gene | (37) |
| *E. coli* |  |  |  |
| SM10(λpir) | ED222 | *thi thr leu tonA lacY supE recA::RP4-2-Tc::Mu Km λpir* | Lab collection |
| χ7213 | ED743 | *thr-1 leuB6 fhuA21 lacY1 glnV44 recA1 ΔasdA4 $\Delta$(zhf-2::Tn10) thi-1 RP4-2-Tc::Mu[$\lambda$ pir]* | Lab collection |

**Table 2.**
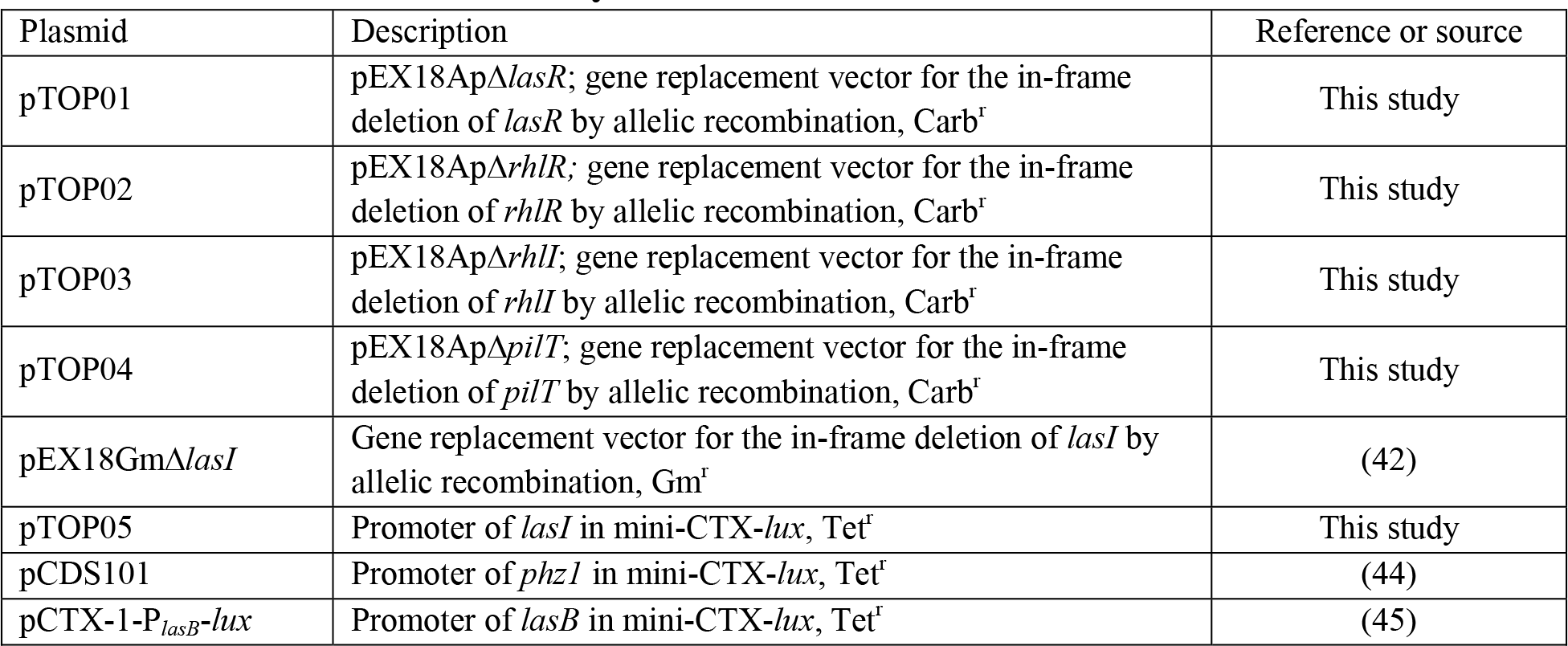
Plasmids used in this study

### Construction of in-frame deletion mutants

An allelic exchange technique based on the use of a suicide vector was used to construct gene knockout deletions (46). Mutant alleles, flanked by regions of homology to the recipient chromosome, were synthesized *in vitro* by PCR from PA14 genomic DNA and then cloned into the allelic exchange vector pEX18Ap (yielding pTOP01, pTOP02, pTOP03, and pTOP04). Plasmids were assembled from purified PCR products and restriction enzyme-cleaved plasmid backbone by employing a seamless strategy of ligation-independent cloning (pEASY® -Uni Seamless Cloning and Assembly Kit, TransGen Biotech Co.). These suicide vectors were transferred into *P. aeruginosa* by conjugation with *E. coli* donor strain (SM10). Carbenicillin was used to select recipient merodiploid cells and *E. coli* donor cells were counter-selected using Irgasan. Double-crossover mutants were isolated by sucrose counter-selection and confirmed by PCR.

### Inactivation of *pilU* gene

Transfer of transposon insertion (::MrT*7*) from the PA14 non-redundant transposon insertion mutant library was used (47) to inactivate *pilU*. Genomic DNA from *pilU*::MrT*7* (mutant ID # 53607) was extracted and transformed into the recipient PA14 Δ*lasR* background. Gentamicin (15 µg/ml) was used to select transformants.

### Construction of chromosomal reporter strains

The promoter region of *lasI* was PCR-amplified from PA14 genomic DNA. pTOP05 (mini-CTX-*lasI-lux*) was constructed by the assembly of the purified PCR product and the enzyme-cleaved mini-CTX-lux backbone (48). pTOP05, pCTX-1-P*_lasB_*-*lux* and pCDS101 were integrated into the *attB* chromosomal site of PA14 and isogenic mutants by conjugation on LB agar plates. Selection was performed on LB agar plates containing tetracycline.

### Luminescence reporter measurements

For *lux* reporter readings, luminescence was measured using a Cytation 3 multimode plate reader (BioTek Instruments, USA). Relative light units (RLU) were normalized by colony-forming units per mL^-1^ (reported in RLU CFU^-1^). When mentioned, AHLs were added to a final concentration of 1.5 μM of C_4_-HSL and 3 μM of 3-oxo-C_12_-HSL from stocks prepared in high-performance liquid chromatography (HPLC)-grade acetonitrile. Acetonitrile only was added in controls.

### Quantification of QS signaling molecules

Concentration of 3-oxo-C_12_-HSL was measured for bacteria grown in liquid King’s A (planktonic growth) and on King’s A agar (surface growth) using HPLC/tandem mass spectrometry (LC/MS/MS) as previously described (49). Quantification was performed at indicated times post-inoculation in both growth conditions. For planktonic growth, overnight cultures grown on TSB were diluted to OD_600_ 0.1 in fresh King’s A medium. At the given time-points, cultures were mixed with acetonitrile containing the internal standard tetradeuterated 4-hydroxy-2-heptylquinoline (HHQ-d_4_), in a 4:1 ratio of culture to solvent (HHQ-d_4_ final concentration of 3 ppm). Bacterial suspension was vortexed and centrifuged at maximum speed for 10 min in order to pellet bacterial cells. The resulting mixture was transferred into vials for LC/MS/MS analyses. Alternatively, for cells grown on agar surfaces, overnight cultures on TSB were diluted to OD_600_ 0.05 in TSB medium. Cultures were grown until an OD_600_ of 1 and agar plugs were inoculated with 5 µl of bacterial suspension. Plates were incubated at 37°C and samples were collected at the indicated time-points. Each sample was composed of two agar plugs mixed with 1 mL of acetonitrile containing the internal standard. This mixture was incubated at 4°C for 16h under gentle agitation, optimizing the diffusion of signaling molecules from the agar to the solvent. After incubation, the mixture was centrifuged at maximum speed for 10 min and the resulting supernatant was transferred into a LC/MS vial. All samples were injected using an HPLC Waters 2795 (Mississauga, ON, Canada) on a Kinetex C8 column (Phenomenex) with an acetonitrile-water gradient containing 1% acetic acid. The detector was a tandem quadrupole mass spectrometer (Quattro premier XE; Waters) equipped with a Z-spray interface using electrospray ionization in positive mode (ESI+). Nitrogen was used as a nebulizing and drying gas at flow rates of 15 and 100 ml · min^−1^, respectively. Concentration was normalized by CFUs per mL^-1^ and reported in ng CFU^-1^. All experiments were performed in triplicates and repeated at least twice independently.

### Pyocyanin quantification

Quantification of pyocyanin produced by surface-grown cells was performed similarly to previously described in (50). Overnight cultures were diluted and grown in TSB until an OD_600_ of 1. At this point, 5 μL were used to inoculate agar plugs from a 96-well plate containing King’s A agar supplemented with FeCl_3_ (200 μL per well). Plates were incubated at 37°C for 24h. Pyocyanin was extracted in 500 μL of chloroform from two agar plugs (by replicate). Tubes were vortexed and centrifuged for 3 min at 12,000 x *g*. 200 μL of the organic phase was recovered in a new tube and a second chloroform extraction was performed on the plugs. The organic phase (400 μL) was acidified with 500 μL 0.2 N HCl and vortexed. The samples were centrifuged for 3 min at 12,000 x *g* and the absorbance of the pink aqueous phase was read at OD_520 nm_. Blank was performed by pyocyanin extraction from uninoculated agar plugs. Values were corrected by colony-forming units per mL^-1^ from samples prepared in the same conditions.

### *Drosophila melanogaster* feeding assay

Fruit flies (*D. melanogaster*) were infected orally in a feeding assay model (51). Male flies (4- to 6-days old) were anesthetized under a gentle stream of carbon dioxide and separated into vials, each containing 10 males. Each strain (or condition) tested was composed of three independent vials, totalizing 30 flies. Vials were prepared with 5 mL of a solution of sucrose agar (5% of sucrose and 1.5% agar). Once solidified, a sterile filter disk was placed on the surface. Prior to infection, bacteria were grown in 6 mL of TSB until an OD_600_ of 3. At this point, the bacterial suspension was centrifuged 3 min at 12,000 x *g* and the pellet was resuspended in 100 μL of sterile 5% sucrose and dispensed on the filter papers. Sterile 5% sucrose alone was used as control. Males were starved 6-8h prior to the infection. Flies were kept at 25°C and about 50% humidity. They were subjected to 12 hrs cycles of light/dark. Mortality was monitored daily for 8 days. The experiment was performed twice, each time in triplicate.

## RESULTS

### Surface growth induces production of 3-oxo-C_12_-HSL in the absence of LasR

In *P. aeruginosa* prototypical strains such as PA14, the quorum sensing regulatory cascade is considered to be primarily activated by the *las* system. LasR, once activated by the binding of 3-oxo-C_12_-HSL, regulates the transcription of target genes, including the gene coding the LasI synthase. This process induces the production of more 3-oxo-C_12_-HSL, resulting in a positive feedback loop. In standard laboratory liquid cultures of *P. aeruginosa*, production of 3-oxo-C_12_-HSL peaks early and decreases overtime ((11); **Fig. 1A**). We note the same pattern of production in wild-type *P. aeruginosa* PA14 (WT) cells grown on an agar surface (**Fig. 1A**). Surprisingly, in a LasR-negative background, the production pattern of 3-oxo-C_12_-HSL is influenced by suspended vs surface culture conditions (**Fig. 1**). As expected, production of the LasR ligand is barely detectable at the stationary phase of a Δ*lasR* mutant in broth cultures. However, its concentration is elevated during surface growth (**Figs. 1A and B**). In WT culture, the peak concentration is observed during the exponential growth phase while it shifts to late-stationary phase in the Δ*lasR* mutant, solely when growing on the surface. This shift might indicate a role for other regulators in the activation of *lasI* transcription in the absence of LasR.

**Figure 1.**
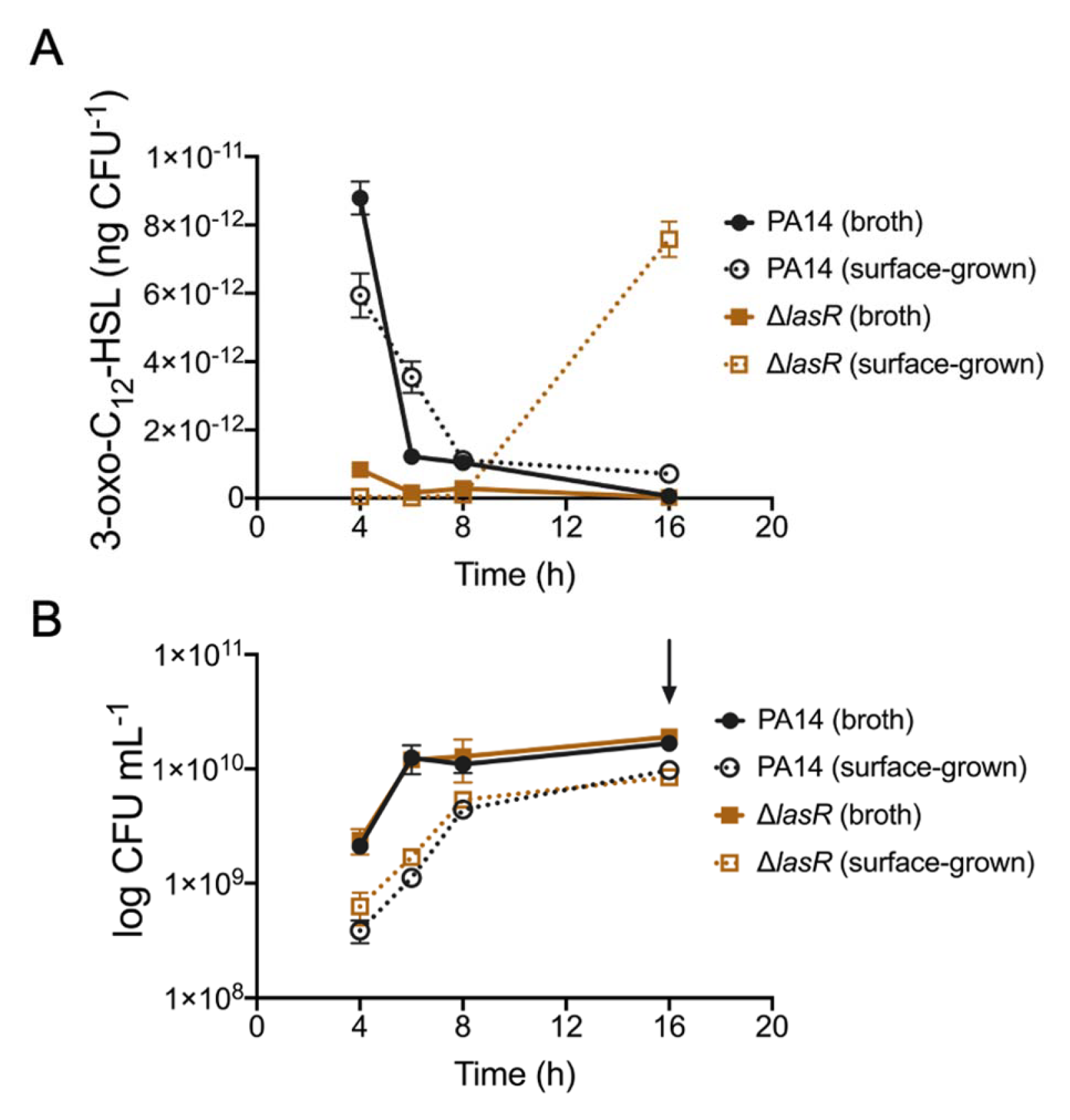
Surface growth induces 3-oxo-C_12_-HSL production in PA14 LasR-null strain. (A) 3-oxo-C_12_-HSL concentration was measured in PA14 and the isogenic Δ*lasR* mutant (PA14 Δ*lasR*) at different time points during planktonic (broth culture) and surface growth (surface of agar-solidified culture media) by liquid chromatography/mass spectrometry. Values were normalized by the viable cell counts and shown in ng CFU^-1^ (B) Growth in broth and surface conditions was determined by the count of viable cells per millilitre (CFU mL^-1^). The arrow indicates the time-point at which 3-oxo-C_12_-HSL is induced in a Δ*lasR* mutant in (A). The values are means ± standard deviation (error bars) from three replicates.

### Production of 3-oxo-C_12_-HSL and expression of *lasI* are RhlR-dependant in LasR-negative backgrounds

Expression of the gene coding the LasI synthase, responsible for the synthesis of 3-oxo-C_12_-HSL, is typically considered to be regulated by LasR. Therefore, little to no production of this AHL is expected in LasR-defective strains, which is what is observed in planktonic cultures. However, upon surface growth, 3-oxo-C_12_-HSL is produced in the absence of LasR. To make sure the production of 3-oxo-C_12_-HSL in this condition still requires LasI activity, we measured concentrations of this AHL in a Δ*lasI* mutant grown under the same surface-associated conditions. As expected, 3-oxo-C_12_-HSL is not detectable in a Δ*lasI* mutant, irrespective of the growth phase (**Fig. 2A**). This result suggests that transcription of *lasI* can occur in absence of LasR upon surface growth. To further investigate this, we measured the activity of a chromosomal *lasI-lux* reporter in a Δ*lasR* background in both planktonic and surface-grown cells. In agreement with the production of 3-oxo-C_12_-HSL, transcription of *lasI* was observed in LasR-negative background grown on a surface (**Fig. 2B**).

**Figure 2.**
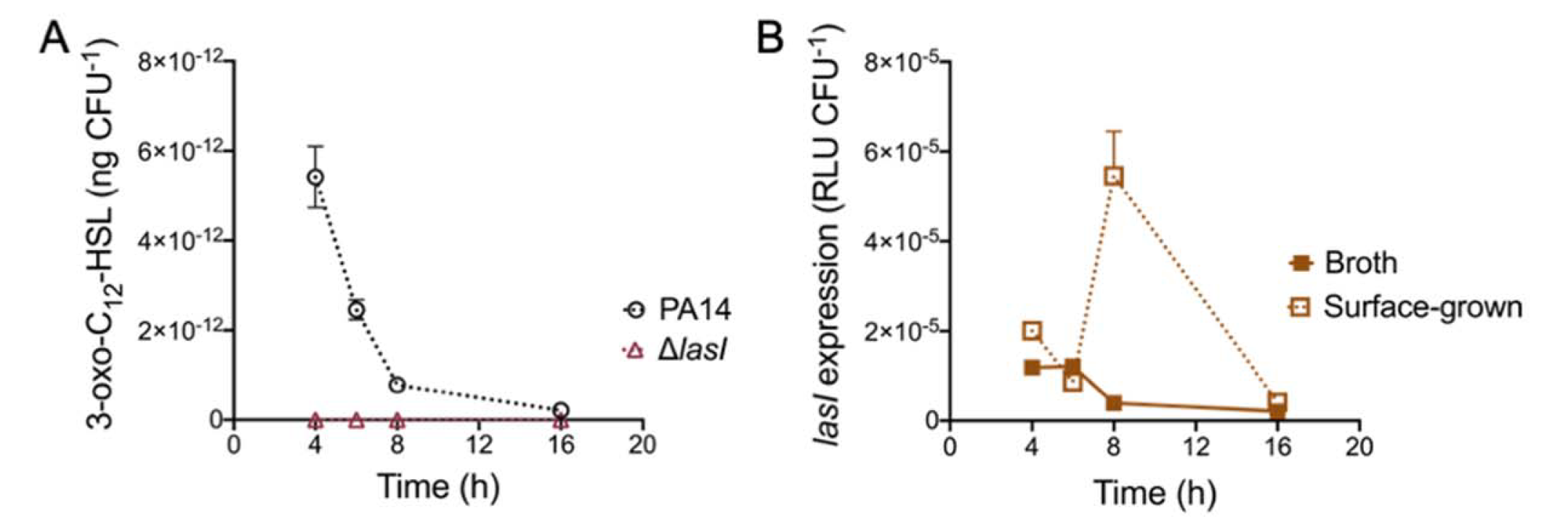
Transcription of *lasI* can occur in absence of LasR in cells growing on a surface. (A) 3-oxo-C_12_-HSL was measured in PA14 and its isogenic Δ*lasI* mutant at different time points during surface growth by liquid chromatography/mass spectrometry. (B) Transcription activity from the chromosomal *lasI*-*lux* reporter in a Δ*lasR* background.

We have previously reported that RhlR can act as a surrogate regulator of LasR-dependent factors in the absence of LasR (38). In *P. aeruginosa* planktonic cultures, this activation is seen by the production of 3-oxo-C_12_-HSL at late stationary phase in LasR-negative backgrounds. However, as shown in **Fig. 1A**, the concentration of this AHL in a Δ*lasR* mutant in broth cultures remains extremely low early on. In contrast, surface growth readily induces production and the corresponding upregulation of *lasI* transcription in a Δ*lasR* mutant (**Figs. 1A and 2B**). To verify if RhlR is responsible for this upregulation, we measured concentrations of 3-oxo-C_12_-HSL in a Δ*rhlR* and a double *lasR rhlR* mutant (**Figs. 3 and S1**) upon surface growth. The production profile of 3-oxo-C_12_-HSL is similar between the WT and a Δ*rhlR* mutant, peaking at exponential growth phase and decaying overtime (**Fig. S1**). The concomitant inactivation of *lasR* and *rhlR* abrogates 3-oxo-C_12_-HSL production, which concurs with our previous finding of RhlR being the alternative activator of *lasI* in LasR-negative backgrounds (**Figs. 3 and S1**). This result suggests that the transcription of *lasI* is mediated by RhlR in surface-grown cells.

**Figure 3.**
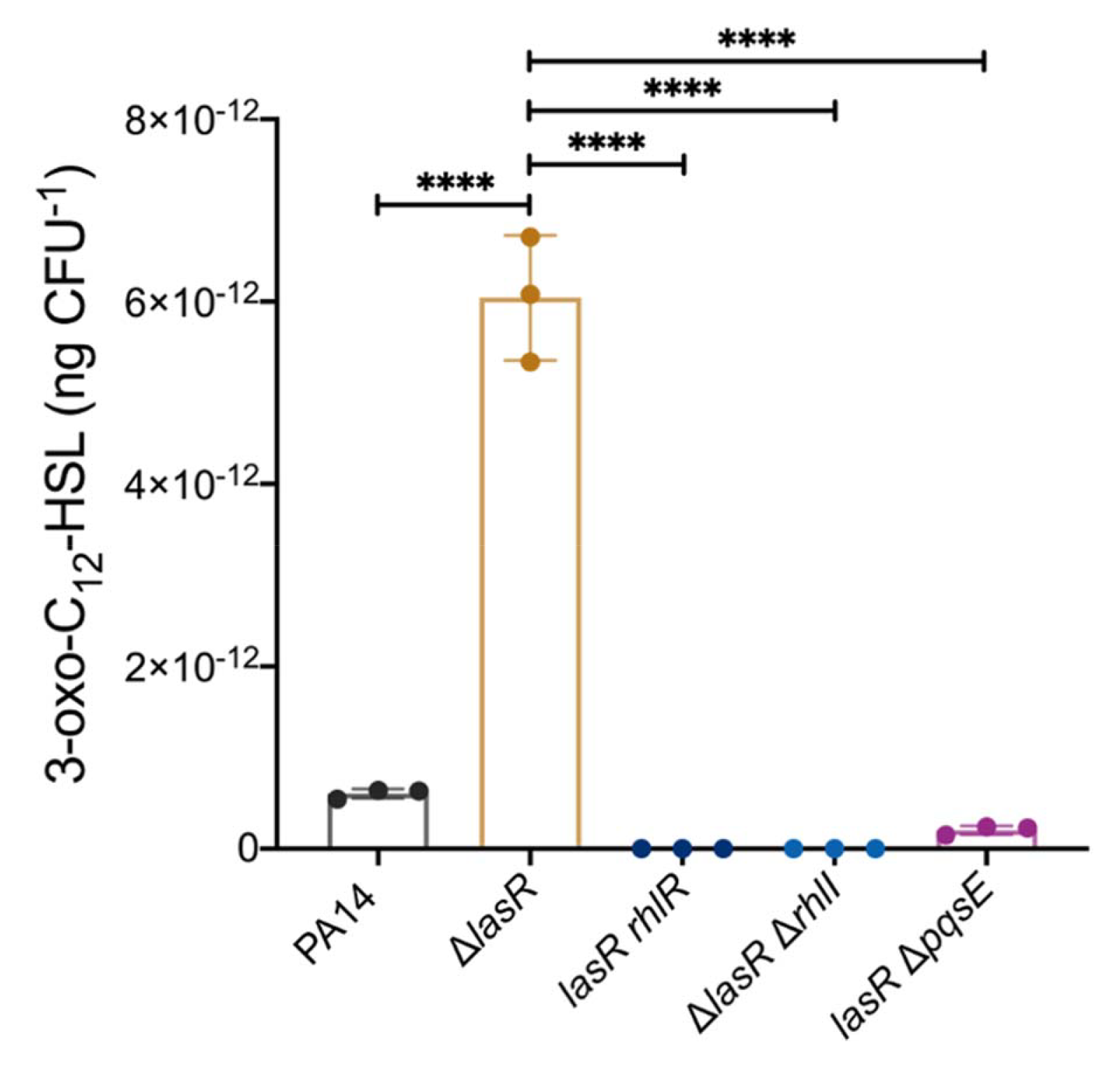
Activity of the Rhl system is required to induce the production of 3-oxo-C_12_-HSL upon surface growth. 3-oxo-C_12_-HSL was measured in PA14, isogenic single-mutants Δ*lasR* and Δ*rhlR*, and the double-mutants *lasRrhlR*, Δ*lasR*Δ*rhlI*, and *lasR*Δ*pqsE* at 16h of surface growth by LC/MS. Concentration was normalized by the viable cell count. The values are means ± standard deviation (error bars) from three replicates. One-way analysis of variance (ANOVA) and Tukey’s multiple comparisons posttest was used to quantify statistical significance. **** *P* ≤ 0.0001.

Thus, in the absence of LasR, surface-grown cells appear to rely on the activity of the *rhl* system to control QS-regulated factors, including the production of 3-oxo-C_12_-HSL. Since the full activity of RhlR depends on both C_4_-HSL and PqsE (13), we measured the concentration of 3-oxo-C_12_-HSL in the double mutants Δ*lasR* Δ*rhlI* and *lasR* Δ*pqsE* in order to further elucidate the role of the Rhl system in this mechanism. As expected, inactivating *rhlI* or *pqsE* in a *lasR* background severely affects the production of 3-oxo-C_12_-HSL (**Fig. 3**) and confirm that the production of 3-oxo-C_12_-HSL by LasR-cells growing on a surface is dependent on the RhlR-mediated transcription of *lasI*.

### Induction of the production of 3-oxo-C_12_-HSL upon surface growth is a widespread response among *P. aeruginosa* strains

Conserved regulation pathways strongly suggest the importance of bacterial responses to their fitness (52). We have observed that surface growth induces production of 3-oxo-C_12_-HSL in an engineered *lasR* deletion mutant of *P. aeruginosa* PA14. To verify if this response is restricted to this prototypical strain, we measured concentrations of this AHL in six naturally occurring LasR-defective *P. aeruginosa* isolates: three strains we recently identified among a collection of environmental isolates (32) and the other three are LasR-defective CF clinical isolates (E41, E113 and E167) from the Early *Pseudomonas* Infection Control (EPIC) study (31, 37). Timing of sampling was chosen based on the 3-oxo-C_12_-HSL production profile of PA14 Δ*lasR*, which peaks at late exponential phase (**Fig. 1**). Considering that growth curves can differ greatly between *P. aeruginosa* strains, we decided to also include a 24h time-point. Environmental and clinical LasR-negative strains behave similarly to the engineered PA14 Δ*lasR* mutant, with production of 3-oxo-C_12_-HSL being augmented upon surface growth when compared to planktonic (**Fig. 4**). The production profile varies among the LasR-negative backgrounds: strain 18G steadily produces 3-oxo-C_12_-HSL during surface growth. At 24h, there is 6-fold more in surface than in the planktonic growth conditions. The environmental strain 32R and the clinical strain E113 have production profiles similar to PA14 Δ*lasR*, and the concentration of 3-oxo-C_12_-HSL peaks at the late exponential phase (**Figs. 4 and S2**). Production is advanced (compared with PA14 Δ*lasR*) in strains 78RV and E167. In these strains, AHL production peaks at early exponential phase (**Figs. 4 and S2**). Finally, upregulation of 3-oxo-C_12_-HSL production upon surface growth was not observed for the clinical strain E41 under our test conditions. Taken together, these results confirm that the absence of a functional LasR generally induces the production of 3-oxo-C_12_-HSL in response to growth in association with surfaces, despite the general requirement of LasR to produce this AHL in standard laboratory planktonic culture conditions.

**Figure 4.**
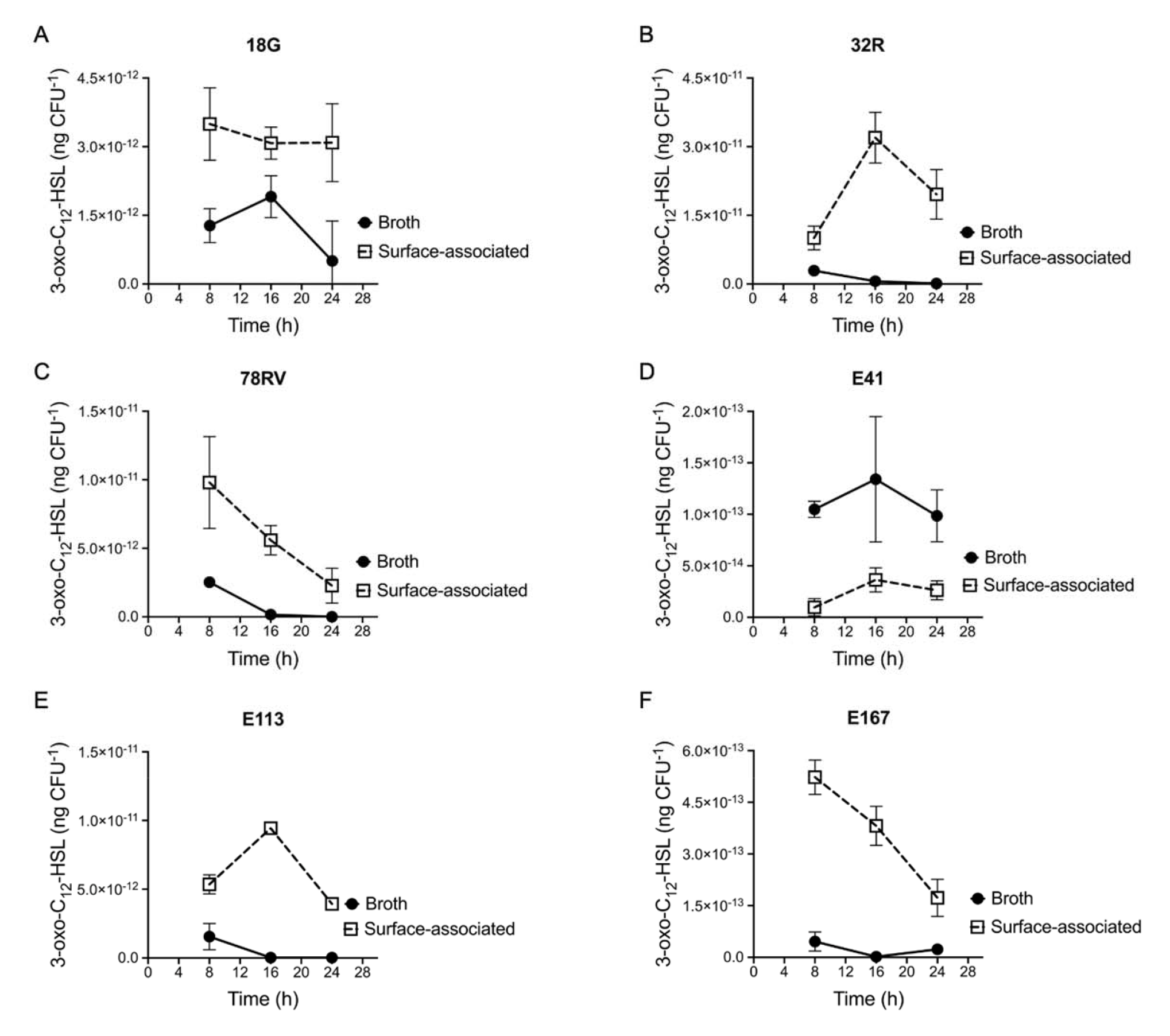
Production of 3-oxo-C_12_-HSL is a widespread feature among LasR-defective strains growing on a surface. 3-oxo-C_12_-HSL was measured at different time-points during planktonic and surface growth by LC/MS of naturally evolved LasR-defective strains. (A) 18G. (B) 32R. (C) 78RV. (D) E41. (E) E113. (F) E167. Concentration was normalized by viable cell count and is shown in ng CFU^-1^. The values are means ± standard deviation (error bars) from three replicates.

### 3-oxo-C_12_-HSL induces the expression of pyocyanin in the absence of LasR

The conservation of surface-primed induction of 3-oxo-C_12_-HSL production in LasR-defective isolates strongly suggests that this signaling molecule mediates significant biological responses in this context. Because 3-oxo-C_12_-HSL is only/essentially known as the autoinducing ligand of LasR, in a LasR-defective background, its production could be considered as a waste of resources. Thus, a plausible explanation for the conservation is that, in the absence of a functional LasR, 3-oxo-C_12_-HSL remains beneficial when *P. aeruginosa* is growing on a surface. Pyocyanin production relies on the expression of the redundant operons *phzA1B1C1D1E1F1G1* (*phz1*) and *phzA2B2C2D2E2F2G2* (*phz2*) – culminating in the synthesis of phenazine-1-carboxylic acid (PCA). PCA is converted to several phenazines, including pyocyanin, the blue pigment characteristic of *P. aeruginosa* cultures (53). Transcription of the *phz1* operon relies on RhlR (13, 54). To verify if 3-oxo-C12-HSL could be implicated in RhlR-dependant QS, we evaluated the level of transcription from the *phz1* promoter during surface-growth, using a chromosomal *phzA1-lux* fusion reporter, in an AHL-and LasR-negative triple mutant (Δ*lasR*Δ*lasI*Δ*rhlI*). As expected, no transcription is seen in the control condition or when only 3-oxo-C_12_-HSL is provided, and upon addition of exogenous C_4_-HSL, *phz1* transcription is induced, consistent with the requirement of C_4_-HSL for RhlR activity (**Fig. 5A**). However, unexpectedly, combined addition of C_4_-HSL and 3-oxo-C_12_-HSL further induces the expression of *phz1* (**Fig. 5A**). The synergetic activation of these signal molecules is also seen for pyocyanin production (**Fig 5B**). The concomitant addition of C_4_-HSL and 3-oxo-C_12_-HSL induces by almost 3-fold the production of this redox-active molecule compared to the addition of C_4_-HSL alone. Similar to the observed *phz1* expression, 3-oxo-C_12_-HSL alone is not sufficient to induce pyocyanin production. These results clearly demonstrates that 3-oxo-C_12_-HSL modulates QS-regulated responses even in the absence its cognate response regulator LasR. This activity depends on the presence of C_4_-HSL, thus likely on the function of RhlR.

**Figure 5.**
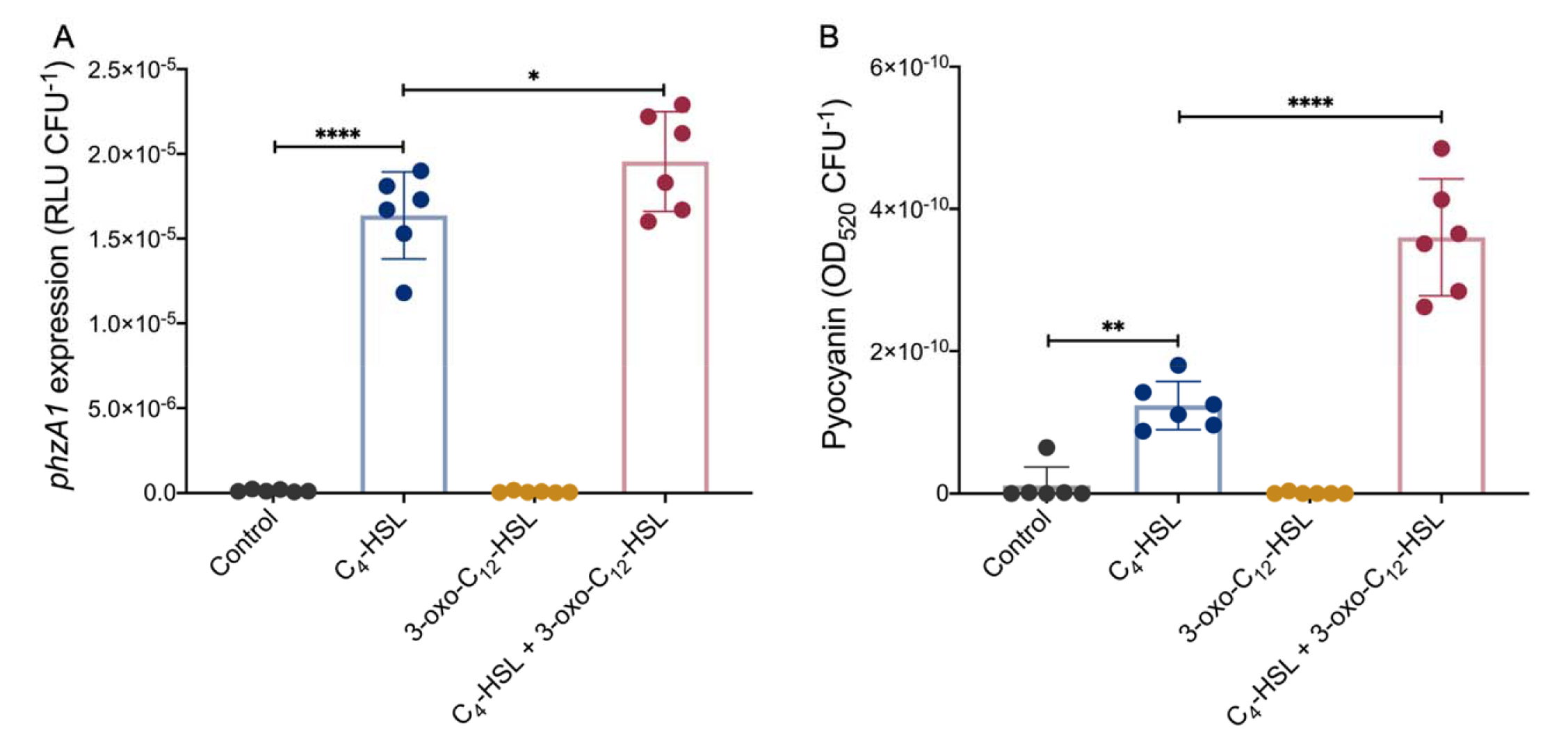
Exogenous 3-oxo-C_12_-HSL induces transcription of the operon *phz1* and pyocyanin production in a *lasR* negative background. (A) Luminescence of the *phzA1-lux* chromosomal reporter was measured in the AHL-negative LasR-defective background (Δ*lasR*Δ*lasI*Δ*rhlI*) after the addition of 1.5 μM of C_4_-HSL, 3 μM of 3-oxo-C_12_-HSL or both molecules at late stationary phase (24h). Acetonitrile alone was used as control. Relative light units were normalized by viable cell counts and shown in RLU CFU^-1^. (B) Pyocyanin produced by Δ*lasR*Δ*lasI*Δ*rhlI* in response to exogenous AHLs was chloroform-extracted at 24h. Production was normalized by cell viable counts and shown in OD_520_ CFU^-1^. The values are means ± standard deviations (error bars) from six replicates. Statistical analyses were performed using ANOVA and Tukey’s multiple comparisons posttest with * *P* ≤ 0.05*; \**\* *P* ≤ 0.01 and **** *P* ≤ 0.0001.

### 3-oxo-C_12_-HSL produced by LasR-negative strains positively regulates the LasB virulence determinant in cocultures

AHLs are conserved extracellular intraspecies signaling molecules. Based on this characteristic, we wondered if 3-oxo-C_12_-HSL produced by LasR-defective isolates could be used by surrounding LasR-active cells to induce LasR-dependent factors. These factors include several exoproducts such as proteases (e.g. LasA and LasB) that can be used by the whole population (“public goods”). To verify this, we measured the activity of the chromosomal *lasB-lux* reporter inserted in Δ*lasI* mutant (Δ*lasI*::CTX *lasB*-*lux* background) in a surface-associated coculture with a Δ*lasR* mutant. Because the Δ*lasI* mutant is unable to produce 3-oxo-C_12_-HSL, the *las* system cannot be activated in this background; however, this strain is LasR-active and prone to induction by exogenous 3-oxo-C_12_-HSL. As expected, *lasB* transcription is at basal levels in Δ*lasI* mutant monoculture (**Fig. 6**). Coculture with Δ*lasR*, which produces 3-oxo-C_12_-HSL under these surface culture conditions, induces the transcription of the *lasB*-lux reporter by more than 4-fold at late stationary phase in which the concentration of LasR-inducing 3-oxo-C_12_-HSL is at its peak. This upregulation depends solely on the production of 3-oxo-C_12_-HSL by the Δ*lasR* mutant, as it is not seen in cocultures with the double mutant Δ*lasR*Δ*lasI*. Thus, 3-oxo-C_12_-HSL produced by LasR-negative strains can be used by surrounding LasR-active cells, modulating the expression of the QS-regulated genes at the communal level.

**Figure 6.**
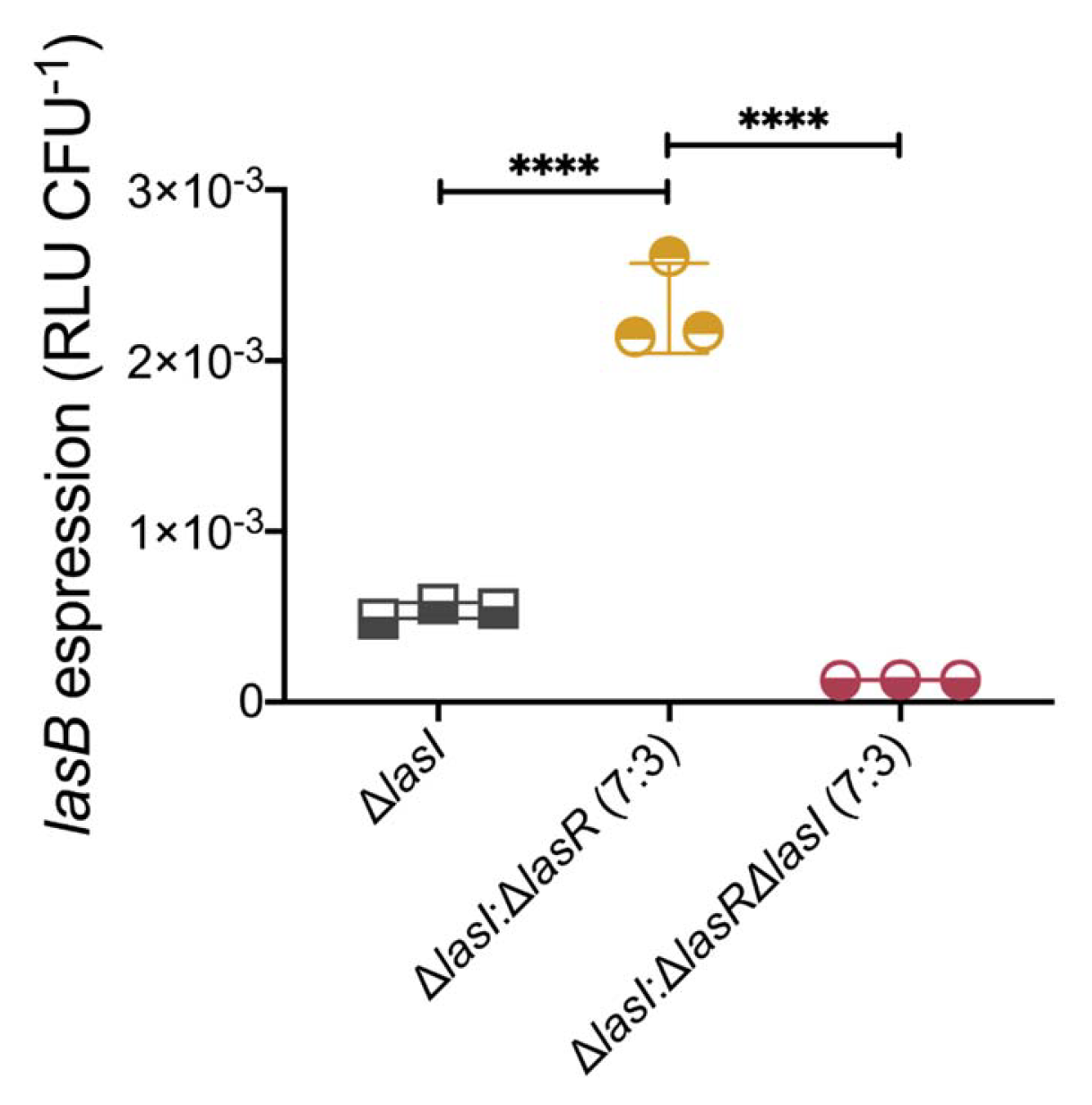
Surface-grown LasR-active cells utilize 3-oxo-C_12_-HSL produced by surrounding LasR-defective mutants, inducing *lasB* expression. Luminescence reading of a *lasB-lux* chromosomal reporter inserted in a LasR-active Δ*lasI* mutant (Δ*lasI*::CTX *lasB-lux*). Monoculture of Δ*lasI* was used as control (gray). Coculture Δ*lasI* and △*lasR* with 7:3 Δ*lasI*-to-Δ*lasR* cell initial ration (yellow). Coculture of Δ*lasI* and Δ*lasR* Δ*lasI* with 7:3 Δ*lasI*-to-Δ*lasR*Δ*lasI* cell initial ratio (red). Relative light unit was normalized by viable cell count of Δ*lasI*::CTX *lasB-lux* strain at 16h and is shown in RLU CFU^-1^. The values are means ± standard deviations (error bars) from three replicates. Statistical significance was calculated by ANOVA and Tukey’s multiple comparisons posttest with **** *P* ≤ 0.0001.

### Virulence of *P. aeruginosa* is positively modulated by LasR-defective cells in coinfection

Even in absence of a functional LasR or endogenous production of its cognate autoinducer, virulence traits are positively regulated by 3-oxo-C_12_-HSL at the individual and community levels (**Figs. 5 and 6**). Thus, we postulated that a coinfection with a mixture of LasR-responsive and LasR-defective strains would be more virulent than a separate infection with the respective strains. To test this, we used the fruit fly *Drosophila melanogaster* as an infection host – in which *P. aeruginosa* cause a disease and mortality (51). We fed fruit flies with *P. aeruginosa* cells and monitored survival of the flies for 8 days post-infection. Feeding assay mimics a chronic infection (55). Virulence of WT PA14 (LasR-active) and Δ*lasR* mutant (LasR-defective) were tested, as well as a coinfection with a 7:3 ratio of each, respectively (**Figs. 7 and S3)**. To address the role of 3-oxo-C_12_-HSL in this response, we also assessed the virulence of the double mutant Δ*lasR*Δ*lasI* in individual and coinfection settings (**Figs. 7 and S3)**. Under our conditions, fly death was accelerated at the beginning by coinfection of PA14 and Δ*lasR*, while overall survival was similar between the later and WT PA14. However, virulence of the coinfection PA14 and Δ*lasR*Δ*lasI* was severely attenuated, indicating that 3-oxo-C_12_-HSL modulates virulence in coinfection settings.

**Figure 7.**
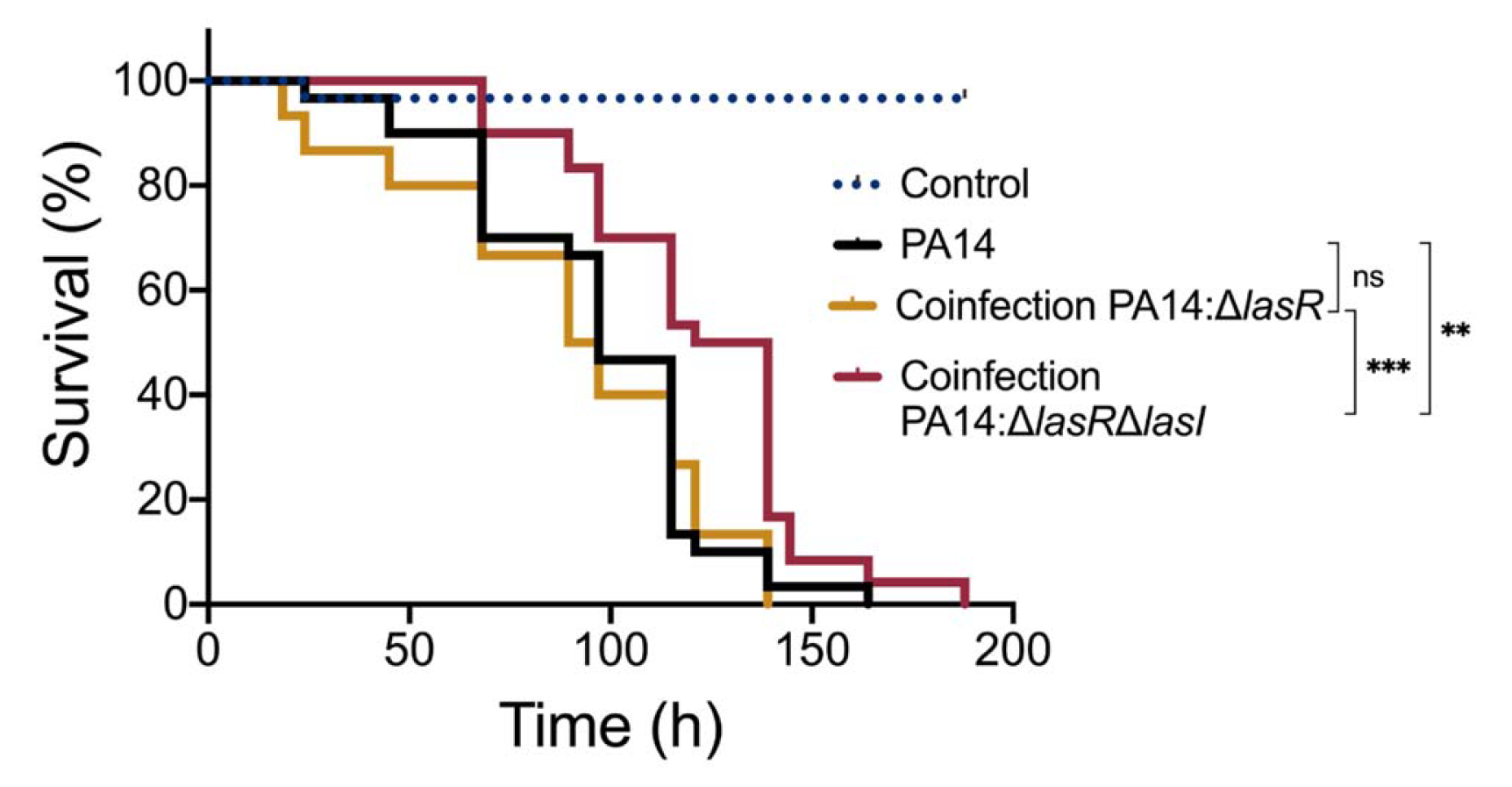
Coinfection of PA14 with the 3-oxo-C_12_-HSL producing Δ*lasR* mutant induces virulence of *P. aeruginosa* toward *D. melanogaster*. Fruit flies were infected with suspended cells in 5% sucrose. Fly survival was monitored over time. *n* = 30 flies per group for each experiment. Experiment was performed independently twice. Statistical significance was determined using Mantel-Cox survival analysis. ns, non-significant, ** *P* ≤ 0.01, and *** *P* ≤ 0.001.

## DISCUSSION

The characteristics and behaviors displayed by bacteria within biofilms have been extensively investigated over the years. These surface-associated communities exhibit features that clearly distinguish them from the free-living counterpart. This is due to a sequential and highly regulated process that mediate the transition from planktonic to sessile lifestyle (56). Although QS regulates social behaviors, often also modulated by aspects related to the sessile way of life, it has been essentially characterized genetically and biochemically in cells grown in broth. In the present study, we show that surface-association is sufficient to induce the LasR-independent expression of *lasI* in *P. aeruginosa* and that 3-oxo-C_12_-HSL modulates the expression of virulence determinants even in the absence of the cognate transcriptional regulator LasR.

Surface-sensing has been previously linked to differential bacterial responses. For instance, we have shown that regulation of the small RNAs RsmY/RsmZ is modulated differently in broth versus surface-grown cells, probably aiding bacterial adaptation to growth conditions (57). Similarly, expression of *lasR* increases in a surface-dependent manner, culminating in a surface-primed QS activation, due to the sensitization of surface-grown cells to the cognate AHL 3-oxo-C_12_-HSL (39). Therefore, QS of *P. aeruginosa* responds differently to the same concentration of 3-oxo-C_12_-HSL: weaker QS activation is seen in planktonic cultures, in contrast to high QS activation in surface-associated cells. This mechanism is reported to rely on type IV (TFP) pili retraction, as surface-primed *lasR* upregulation is lost in the absence of the motors PilT and PilU (39). Thus, a relationship between QS and surface sensing is established but its complexity remains to be clearly defined.

LasR-defective *P. aeruginosa* isolates have been generally related to human chronic infections, in which this bacterium persists in the lungs of people with CF as a biofilm. Recently, the general high occurrence of such isolates challenged this long-held notion (32, 33). Loss of LasR function appears to be a widespread adaptation feature of this bacterium (32). Our results support a model in which surface attachment, a growth condition often encountered by *P. aeruginosa*, induces RhlR-dependent production of 3-oxo-C_12_-HSL in LasR-defective background – sustaining QS-responsiveness in this condition. Why is RhlR-dependent expression of LasI observed in a LasR-deficient background more prominent in sessile cells? Compared to planktonic growth, both sessile LasR-active and LasR-defective cells produce more C_4_-HSL (**Fig. S4**), which could lead to a stronger activation of the *rhl* system, culminating in the upregulation of RhlR-dependent factors. However, the RhlR-dependent 3-oxo-C_12_-HSL overproduction in sessile cells is seen only in LasR-defective backgrounds.

Irrespective of the mechanism, the production of 3-oxo-C_12_-HSL appears to have important biological implications. As mentioned before, surface-association upregulates LasR, thus sensitizing cells to 3-oxo-C_12_-HSL (39). Upregulation of 3-oxo-C_12_-HSL in LasR-defective background does not appear dependent on this mechanism, as the TFP retraction motors PilT and PilU are not required for this response (**Fig. S5**).

The conservation of surface-primed induction of 3-oxo-C_12_-HSL in naturally occurring LasR-defective isolates of *P. aeruginosa* is an indicator of its importance. We observed this response in naturally-evolved LasR-defective isolates from both clinical and environmental origins (32, 43). The environmental isolates used here, namely 18G, 32R and 78RV were recently characterized as LasR-defective strains based on their inability to perform LasR-dependent activities in liquid cultures (32). Of note, due to the ability to mediate RhlR-regulated QS, LasR-defective 78RV was characterized as a RAIL strain (32), like the CF isolates E113 and E167, which also have functional RhlR-dependent QS responses (37). In the absence of a functional LasR, surface-association induces the production of the 3-oxo-C_12_-HSL signal irrespective of the QS-responsiveness mediated by LasR-independent RhlR. This response is prevalent, but not universal. Isolate E41 produces trace concentrations of 3-oxo-C_12_-HSL and its production was not induced by surface-association when compared with planktonic cells. Response variability is not surprising considering the diversity of *P. aeruginosa* isolates, but our results highlight that LasR-deficient *P. aeruginosa* isolated from both clinical and environmental settings are often proficient in the production of 3-oxo-C_12_-HSL when adopting an attached growth mode. Thus, this ability appears to be an intrinsic and beneficial feature of this species.

Mutations in the cognate synthase gene, *lasI*, are much less frequently detected than those found in the *lasR* gene (33). The most accepted explanation for this discrepancy is social cheating. Cheaters are individuals that benefit of a shared beneficial product or function (“public good”) while contributing less than average to the metabolic cost. Inactivation of LasI would not prevent response to 3-oxo-C_12_-HSL produced by neighboring WT cells and thus activate a functional LasR. LasR-defective isolates emerge even in experimental conditions that do not apparently require QS-induced products (and thus cheating) (58). An alternative explanation for a lower frequency of *lasI*-null isolates is that 3-oxo-C_12_-HSL might contribute an alternative function beyond LasR activation. This interpretation is supported by our results, where LasR-defective strains retain the ability to respond to the presence of 3-oxo-C_12_-HSL. Indeed, expression of *phz1*, a QS-regulated operon required for pyocyanin production, is controlled by RhlR and its cognate ligand C_4_-HSL. Concomitant addition of 3-oxo-C_12_-HSL further induces *phz1* transcription, and positively regulates pyocyanin production suggesting a response to this non-cognate AHL (**Fig. 5**). The induction of RhlR-controlled *phz1* expression by 3-oxo-C_12_-HSL is also seen in the double mutant Δ*lasR*Δ*lasI* (**Fig. S6**). Basal expression of *phz1* is due to the self-produced C_4_-HSL. Addition of 3-oxo-C_12_-HSL further enhances *phz1* transcription activity and the highest expression is seen when C_4_-HSL is added with 3-oxo-C_12_-HSL. The RhlR-dependent response to the non-cognate signal 3-oxo-C_12_-HSL remains to be understood, but further supports the importance of maintaining LasI activity in the absence of LasR.

Producing 3-oxo-C_12_-HSL in the absence of LasR can also have a positive community outcome. Because it is exported, we have shown that this AHL can have exogenous effects in surrounding cells in a surface-associated setting. Thus, localized production of 3-oxo-C_12_-HSL by LasR-negative clusters could induce the expression of QS-regulated virulence factors in LasR-active cells, with minimal metabolic cost to the LasR-negative producers. Moreover, the production profile is delayed in LasR-defective strains when compared to the WT. Therefore, the mixed population composed of both LasR-active and LasR-defective cells would be subjected to steady levels of 3-oxo-C_12_-HSL. Furthermore, in natural habitats, *P. aeruginosa* is part of complex polymicrobial communities. Microbes within these communities can actively respond to one another, and these interactions range from cooperation to competition (59). Steady production of 3-oxo-C_12_-HSL by *P. aeruginosa* might be relevant in shaping the biological activities of the population. Accordingly, LuxR homologues BtaR1 and BtaR2 – from *Burkholderia thailandensis* – are promiscuous and prone to activation by 3-oxo-C_12_-HSL (60). Importantly, *P. aeruginosa* and *B. thailandensis* are soil saprophytes and thus, may occupy the same environmental niches (61). The ecological relevance of sensing signals produced by neighbouring cells is seen by the conservation of an orphan LuxR homologue (SdiA) in *Salmonella enterica* serovar Typhimurium, a bacterium unable to produce AHLs (62). This ability to “eavesdrop” the environment, by sensing AHL produced by other bacteria is likely not unique in this bacterium and might modulate interspecies interactions and justify the benefit of sustained production of 3-oxo-C_12_-HSL in *P. aeruginosa*. This idea is further supported by the marine sponge symbiont *Ruegeria* sp, a bacterium with a solo LuxI homologue, thus unable to utilize this molecule in a traditional QS-regulated pathway (63).

QS signals also play a pivotal role in host-pathogen interactions. QS-regulated molecules can act as interkingdom QS signals, thus responsible for the communication of bacteria with mammalian cells and the modulation of host immune systems. Indeed, this was reported for 3-oxo-C_12_-HSL (recently reviewed by (64)). Due to the long acyl chain of this autoinducer, the molecule has lipophilic properties and, by directly interacting with biological membranes, can enter mammalian cells and directly interact with intracellular molecules (65). Presence of 3-oxo-C_12_-HSL induces apoptosis of haematopoietic cells and cytotoxicity of non-haematopoietic cells, including those of the airway epithelium (66-70). The host immune responses are also suppressed by 3-oxo-C_12_-HSL, negatively impacting cytokines production, T cell differentiation as well as the function of antigen-presenting cells (71-73). Thus, this signal molecule is central to virulence and pathogenesis of *P. aeruginosa* and the sustained production of 3-oxo-C_12_-HSL by biofilm-growing LasR-deficient isolates in infected hosts might account for worse clinical outcomes. In infected hosts, could the immunomodulatory activity of 3-oxo-C_12_-HSL, rather than its role as quorum sensing signal, justify the regulatory by-pass in the absence of LasR?

Sustained production of 3-oxo-C_12_-HSL in the absence of LasR in response to surface growth, the most common lifestyle adopted by *P. aeruginosa* in its natural environments, appears to be beneficial to the colonization of many environmental niches. Combined with the widespread feature underlying the emergence of LasR-defective isolates, it raises an important question: do these isolates emerge solely to benefit from the cooperating individuals or could they play a positive role in shaping the bacterial community responses?

## ACKNOWLEDGMENTS

We thank George A. O’Toole (Dartmouth) and Matthew T. Cabeen (Oklahoma State University) for gifts of strains and plasmids used in this work. TOP was the recipient of PhD scholarships from the Fondation Armand-Frappier. This research was supported by Canadian Institutes of Health Research (CIHR) operating grant MOP-142466.

## REFERENCES

1. Fuqua WC, Winans SC, Greenberg EP. 1994. Quorum sensing in bacteria: the LuxR-LuxI family of cell density-responsive transcriptional regulators. J Bacteriol 176:269–275.

2. Azimi S, Klementiev AD, Whiteley M, Diggle SP. 2020. Bacterial quorum sensing during infection. Annu Rev Microbiol 74:201–219.

3. Pearson JP, Gray KM, Passador L, Tucker KD, Eberhard A, Iglewski BH, Greenberg EP. 1994. Structure of the autoinducer required for expression of *Pseudomonas aeruginosa* virulence genes. Proceedings of the National Academy of Sciences 91:197–201.

4. Pearson JP, Passador L, Iglewski BH, Greenberg EP. 1995. A second *N*-acylhomoserine lactone signal produced by *Pseudomonas aeruginosa*. Proceedings of the National Academy of Sciences 92:1490–1494.

5. Gambello MJ, Iglewski BH. 1991. Cloning and characterization of the *Pseudomonas aeruginosa lasR* gene, a transcriptional activator of elastase expression. J Bacteriol 173:3000–3009.

6. Schuster M, Urbanowski ML, Greenberg EP. 2004. Promoter specificity in *Pseudomonas aeruginosa* quorum sensing revealed by DNA binding of purified LasR. Proceedings of the National Academy of Sciences 101:15833–15839.

7. Seed PC, Passador L, Iglewski BH. 1995. Activation of the *Pseudomonas aeruginosa lasI* gene by LasR and the *Pseudomonas* autoinducer PAI: an autoinduction regulatory hierarchy. Journal of Bacteriology 177:654–659.

8. Cao H, Krishnan G, Goumnerov B, Tsongalis J, Tompkins R, Rahme LG. 2001. A quorum sensing-associated virulence gene of *Pseudomonas aeruginosa* encodes a LysR-like transcription regulator with a unique self-regulatory mechanism. Proc Natl Acad Sci U S A 98:14613–14618.

9. Gallagher LA, McKnight SL, Kuznetsova MS, Pesci EC, Manoil C. 2002. Functions required for extracellular quinolone signaling by *Pseudomonas aeruginosa*. J Bacteriol 184:6472–6480.

10. Déziel E, Lépine F, Milot S, He J, Mindrinos MN, Tompkins RG, Rahme LG. 2004. Analysis of *Pseudomonas aeruginosa* 4-hydroxy-2-alkylquinolines (HAQs) reveals a role for 4-hydroxy-2-heptylquinoline in cell-to-cell communication. Proc Natl Acad Sci U S A 101:1339–1344.

11. Déziel E, Gopalan S, Tampakaki AP, Lépine F, Padfield KE, Saucier M, Xiao G, Rahme LG. 2005. The contribution of MvfR to *Pseudomonas aeruginosa* pathogenesis and quorum sensing circuitry regulation: multiple quorum sensing-regulated genes are modulated without affecting *lasRI*, *rhlRI* or the production of *N*-acyl-L-homoserine lactones. Mol Microbiol 55:998–1014.

12. Farrow JM, 3rd, Sund ZM, Ellison ML, Wade DS, Coleman JP, Pesci EC. 2008. PqsE functions independently of PqsR-*Pseudomonas* quinolone signal and enhances the rhl quorum-sensing system. J Bacteriol 190:7043-7051.

13. Groleau M-C, de Oliveira Pereira T, Dekimpe V, Déziel E. 2020. PqsE Is essential for RhlR-dependent quorum sensing regulation in *Pseudomonas aeruginosa*. mSystems 5:e00194-20.

14. Letizia M, Mellini M, Fortuna A, Visca P, Imperi F, Leoni L, Rampioni G. 2022. PqsE expands and differentially modulates the RhlR quorum sensing regulon in *Pseudomonas aeruginosa*. Microbiol Spectr 10:e0096122.

15. Wade DS, Calfee MW, Rocha ER, Ling EA, Engstrom E, Coleman JP, Pesci EC. 2005. Regulation of *Pseudomonas* quinolone signal synthesis in *Pseudomonas aeruginosa*. Journal of Bacteriology 187:4372–4380.

16. Xiao G, Déziel E, He J, Lépine F, Lesic B, Castonguay M-H, Milot S, Tampakaki AP, Stachel SE, Rahme LG. 2006. MvfR, a key Pseudomonas aeruginosa pathogenicity LTTR-class regulatory protein, has dual ligands. Molecular Microbiology 62:1689–1699.

17. de Kievit TR, Kakai Y, Register JK, Pesci EC, Iglewski BH. 2002. Role of the *Pseudomonas aeruginosa las* and *rhl* quorum-sensing systems in *rhlI* regulation. FEMS Microbiology Letters 212:101–106.

18. Pesci EC, Pearson JP, Seed PC, Iglewski BH. 1997. Regulation of *las* and *rhl* quorum sensing in *Pseudomonas aeruginosa*. Journal of Bacteriology 179:3127–3132.

19. Mattick JS. 2002. Type IV pili and twitching motility. Annu Rev Microbiol 56:289–314.

20. O’Toole GA, Wong GC. 2016. Sensational biofilms: surface sensing in bacteria. Current Opinion in Microbiology 30:139–146.

21. Kearns DB. 2010. A field guide to bacterial swarming motility. Nature Reviews Microbiology 8:634–644.

22. Galán JE, Collmer A. 1999. Type III secretion machines: bacterial devices for protein delivery into host cells. Science 284:1322–1328.

23. Siryaporn A, Kuchma SL, O’Toole GA, Gitai Z. 2014. Surface attachment induces *Pseudomonas aeruginosa* virulence. Proceedings of the National Academy of Sciences 111:16860–16865.

24. Persat A, Inclan YF, Engel JN, Stone HA, Gitai Z. 2015. Type IV pili mechanochemically regulate virulence factors in *Pseudomonas aeruginosa*. Proceedings of the National Academy of Sciences 112:7563–7568.

25. Williams BJ, Dehnbostel J, Blackwell TS. 2010. *Pseudomonas aeruginosa*: Host defence in lung diseases. Respirology 15:1037–1056.

26. Alhede M, Bjarnsholt T, Givskov M, Alhede M. 2014. *Pseudomonas aeruginosa* biofilms: mechanisms of immune evasion. Adv Appl Microbiol 86:1–40.

27. Moreau-Marquis S, Stanton BA, O’Toole GA. 2008. *Pseudomonas aeruginosa* biofilm formation in the cystic fibrosis airway. Pulm Pharmacol Ther 21:595–599.

28. Smith EE, Buckley DG, Wu Z, Saenphimmachak C, Hoffman LR, D’Argenio DA, Miller SI, Ramsey BW, Speert DP, Moskowitz SM, Burns JL, Kaul R, Olson MV. 2006. Genetic adaptation by *Pseudomonas aeruginosa* to the airways of cystic fibrosis patients. Proceedings of the National Academy of Sciences 103:8487–8492.

29. D’Argenio DA, Wu M, Hoffman LR, Kulasekara HD, Déziel E, Smith EE, Nguyen H, Ernst RK, Larson Freeman TJ, Spencer DH, Brittnacher M, Hayden HS, Selgrade S, Klausen M, Goodlett DR, Burns JL, Ramsey BW, Miller SI. 2007. Growth phenotypes of *Pseudomonas aeruginosa lasR* mutants adapted to the airways of cystic fibrosis patients. Mol Microbiol 64:512–533.

30. Hoffman LR, Kulasekara HD, Emerson J, Houston LS, Burns JL, Ramsey BW, Miller SI. 2009. *Pseudomonas aeruginosa lasR* mutants are associated with cystic fibrosis lung disease progression. J Cyst Fibros 8:66–70.

31. Feltner J, Wolter D, Pope C, Groleau M, Smalley N, Greenberg E, Mayer-Hamblett N, Burns J, Déziel E, Hoffman L, Dandekar A. 2016. LasR variant cystic fibrosis isolates reveal an adaptable quorum-sensing hierarchy in *Pseudomonas aeruginosa*. mBio 7:e01513–16.

32. Groleau MC, Taillefer H, Vincent AT, Constant P, Déziel E. 2022. *Pseudomonas aeruginosa* isolates defective in function of the LasR quorum sensing regulator are frequent in diverse environmental niches. Environ Microbiol 24:1062–1075.

33. O’Connor K, Zhao CY, Mei M, Diggle SP. 2022. Frequency of quorum-sensing mutations in *Pseudomonas aeruginosa* strains isolated from different environments. Microbiology (Reading) 168.

34. Chen R, Déziel E, Groleau M-C, Schaefer AL, Greenberg EP. 2019. Social cheating in a *Pseudomonas aeruginosa* quorum-sensing variant. Proceedings of the National Academy of Sciences 116:7021–7026.

35. Kostylev M, Kim DY, Smalley NE, Salukhe I, Greenberg EP, Dandekar AA. 2019. Evolution of the *Pseudomonas aeruginosa* quorum-sensing hierarchy. Proceedings of the National Academy of Sciences 116:7027–7032.

36. Cruz R, Asfahl K, Van den Bossche S, Coenye T, Crabbé A, Dandekar A. 2020. RhlR-regulated acyl-homoserine lactone quorum sensing in a cystic fibrosis isolate of *Pseudomonas aeruginosa*. mBio 11:e00532–20.

37. Asfahl KL, Smalley NE, Chang AP, Dandekar AA. 2022. Genetic and transcriptomic characteristics of RhlR-dependent quorum sensing in cystic fibrosis isolates of *Pseudomonas aeruginosa*. mSystems 7:e0011322.

38. Dekimpe V, Déziel E. 2009. Revisiting the quorum-sensing hierarchy in *Pseudomonas aeruginosa*: the transcriptional regulator RhlR regulates LasR-specific factors. Microbiology (Reading) 155:712–723.

39. Chuang SK, Vrla GD, Fröhlich KS, Gitai Z. 2019. Surface association sensitizes *Pseudomonas aeruginosa* to quorum sensing. Nature Communications 10:4118.

40. King EO, Ward MK, Raney DE. 1954. Two simple media for the demonstration of pyocyanin and fluorescin. J Lab Clin Med 44:301–307.

41. Rahme LG, Stevens EJ, Wolfort SF, Shao J, Tompkins RG, Ausubel FM. 1995. Common virulence factors for bacterial pathogenicity in plants and animals. Science 268:1899–1902.

42. Hogan DA, Vik A, Kolter R. 2004. A *Pseudomonas aeruginosa* quorum-sensing molecule influences *Candida albicans* morphology. Mol Microbiol 54:1212–1223.

43. Déziel E, Paquette G, Villemur R, Lepine F, Bisaillon J. 1996. Biosurfactant production by a soil *Pseudomonas* strain growing on polycyclic aromatic hydrocarbons. Applied and Environmental Microbiology 62:1908–1912.

44. Sibley CD, Duan K, Fischer C, Parkins MD, Storey DG, Rabin HR, Surette MG. 2008. Discerning the complexity of community interactions using a *Drosophila* model of polymicrobial infections. PLoS Pathogens 4:e1000184.

45. Cabeen MT. 2014. Stationary phase-specific virulence factor overproduction by a *lasR* mutant of *Pseudomonas aeruginosa*. PLoS ONE 9:e88743.

46. Hmelo LR, Borlee BR, Almblad H, Love ME, Randall TE, Tseng BS, Lin C, Irie Y, Storek KM, Yang JJ, Siehnel RJ, Howell PL, Singh PK, Tolker-Nielsen T, Parsek MR, Schweizer HP, Harrison JJ. 2015. Precision-engineering the *Pseudomonas aeruginosa* genome with two-step allelic exchange. Nature Protocols 10:1820–1841.

47. Liberati NT, Urbach JM, Miyata S, Lee DG, Drenkard E, Wu G, Villanueva J, Wei T, Ausubel FM. 2006. An ordered, nonredundant library of *Pseudomonas aeruginosa* strain PA14 transposon insertion mutants. Proc Natl Acad Sci U S A 103:2833–2838.

48. Becher A, Schweizer HP. 2000. Integration-proficient *Pseudomonas aeruginosa* vectors for isolation of single-copy chromosomal *lacZ* and *lux* gene fusions. Biotechniques 29:948–950, 952.

49. Lépine F, Milot S, Groleau M-C, Déziel E. 2018. Liquid chromatography/mass spectrometry (LC/MS) for the detection and quantification of N-acyl-L-homoserine lactones (AHLs) and 4-hydroxy-2-alkylquinolines (HAQs). doi:10.1007/978-1-4939-7309-5_4:49-59.

50. Mould Dallas L, Botelho Nico J, Hogan Deborah A. 2020. Intraspecies signaling between common variants of *Pseudomonas aeruginosa* increases production of quorum-sensing-controlled virulence factors. mBio 11:e01865–20.

51. Apidianakis Y, Rahme LG. 2009. *Drosophila melanogaster* as a model host for studying *Pseudomonas aeruginosa infection*. Nat Protoc 4:1285–1294.

52. Jordan IK, Rogozin IB, Wolf YI, Koonin EV. 2002. Essential genes are more evolutionarily conserved than are nonessential genes in bacteria. Genome Res 12:962–968.

53. Mavrodi DV, Bonsall RF, Delaney SM, Soule MJ, Phillips G, Thomashow LS. 2001. Functional analysis of genes for biosynthesis of pyocyanin and phenazine-1-carboxamide from *Pseudomonas aeruginosa* PAO1. J Bacteriol 183:6454–6465.

54. Whiteley M, Greenberg EP. 2001. Promoter specificity elements in *Pseudomonas aeruginosa* quorum-sensing-controlled genes. Journal of Bacteriology 183:5529–5534.

55. Purdy AE, Watnick PI. 2011. Spatially selective colonization of the arthropod intestine through activation of *Vibrio cholerae* biofilm formation. Proceedings of the National Academy of Sciences 108:19737–19742.

56. Rumbaugh KP, Sauer K. 2020. Biofilm dispersion. Nature Reviews Microbiology 18:571–586.

57. Jean-Pierre F, Tremblay J, Déziel E. 2017. Broth versus surface-grown cells: differential regulation of RsmY/Z small RNAs in *Pseudomonas aeruginosa* by the Gac/HptB system. Front Microbiol 7:2168.

58. Mould DL, Stevanovic M, Ashare A, Schultz D, Hogan DA. 2022. Metabolic basis for the evolution of a common pathogenic *Pseudomonas aeruginosa* variant. eLife 11:e76555.

59. Mitri S, Richard Foster K. 2013. The genotypic view of social interactions in microbial communities. Annual Review of Genetics 47:247–273.

60. Wellington S, Greenberg EP. 2019. Quorum sensing signal selectivity and the potential for interspecies cross talk. mBio 10:e00146–19.

61. Zhao J, Schloss PD, Kalikin LM, Carmody LA, Foster BK, Petrosino JF, Cavalcoli JD, Vandevanter DR, Murray S, Li JZ, Young VB, Lipuma JJ. 2012. Decade-long bacterial community dynamics in cystic fibrosis airways. Proceedings of the National Academy of Sciences 109:5809–5814.

62. Dyszel JL, Smith JN, Lucas DE, Soares JA, Swearingen MC, Vross MA, Young GM, Ahmer BM. 2010. *Salmonella enterica* serovar Typhimurium can detect acyl homoserine lactone production by *Yersinia enterocolitica* in mice. J Bacteriol 192:29–37.

63. Zan J, Choi O, Meharena H, Uhlson CL, Churchill MEA, Hill RT, Fuqua C. 2015. A solo luxI-type gene directs acylhomoserine lactone synthesis and contributes to motility control in the marine sponge symbiont Ruegeria sp. KLH11. Microbiology 161:50-56.

64. Fan Q, Wang H, Mao C, Li J, Zhang X, Grenier D, Yi L, Wang Y. 2022. Structure and signal regulation mechanism of interspecies and interkingdom quorum sensing system receptors. Journal of Agricultural and Food Chemistry 70:429–445.

65. Ritchie AJ, Whittall C, Lazenby JJ, Chhabra SR, Pritchard DI, Cooley MA. 2007. The immunomodulatory *Pseudomonas aeruginosa* signalling molecule *N*-(3-oxododecanoyl)-L-homoserine lactone enters mammalian cells in an unregulated fashion. Immunol Cell Biol 85:596–602.

66. Tateda K, Ishii Y, Horikawa M, Matsumoto T, Miyairi S, Pechere Jean C, Standiford Theodore J, Ishiguro M, Yamaguchi K. 2003. The *Pseudomonas aeruginosa* autoinducer *N*-3-oxododecanoyl homoserine lactone accelerates apoptosis in macrophages and neutrophils. Infection and Immunity 71:5785–5793.

67. Li H, Wang L, Ye L, Mao Y, Xie X, Xia C, Chen J, Lu Z, Song J. 2009. Influence of *Pseudomonas aeruginosa* quorum sensing signal molecule *N*-(3-oxododecanoyl) homoserine lactone on mast cells. Med Microbiol Immunol 198:113–121.

68. Schwarzer C, Fu Z, Patanwala M, Hum L, Lopez-Guzman M, Illek B, Kong W, Lynch SV, Machen TE. 2012. *Pseudomonas aeruginosa* biofilm-associated homoserine lactone C12 rapidly activates apoptosis in airway epithelia. Cell Microbiol 14:698–709.

69. Shiner EK, Terentyev D, Bryan A, Sennoune S, Martinez-Zaguilan R, Li G, Gyorke S, Williams SC, Rumbaugh KP. 2006. *Pseudomonas aeruginosa* autoinducer modulates host cell responses through calcium signalling. Cell Microbiol 8:1601–1610.

70. Kravchenko VV, Kaufmann GF, Mathison JC, Scott DA, Katz AZ, Wood MR, Brogan AP, Lehmann M, Mee JM, Iwata K, Pan Q, Fearns C, Knaus UG, Meijler MM, Janda KD, Ulevitch RJ. 2006. *N*-(3-oxo-acyl)homoserine lactones signal cell activation through a mechanism distinct from the canonical pathogen-associated molecular pattern recognition receptor pathways. J Biol Chem 281:28822–28830.

71. Ritchie AJ, Jansson A, Stallberg J, Nilsson P, Lysaght P, Cooley MA. 2005. The *Pseudomonas aeruginosa* quorum-sensing molecule *N*-3-(oxododecanoyl)-L-homoserine lactone inhibits T-cell differentiation and cytokine production by a mechanism involving an early step in T-cell activation. Infect Immun 73:1648–1655.

72. Telford G, Wheeler D, Williams P, Tomkins PT, Appleby P, Sewell H, Stewart GS, Bycroft BW, Pritchard DI. 1998. The *Pseudomonas aeruginosa* quorum-sensing signal molecule *N*-(3-oxododecanoyl)-L-homoserine lactone has immunomodulatory activity. Infect Immun 66:36–42.

73. Li Y, Zhou H, Zhang Y, Chen C, Huang B, Qu P, Zeng J, Shunmei E, Zhang X, Liu J. 2015. *N*-3-(oxododecanoyl)-L-homoserine lactone promotes the induction of regulatory T-cells by preventing human dendritic cell maturation. Exp Biol Med (Maywood) 240:896–903.

