## Supplemental Figures for "Surface growth of *Pseudomonas aeruginosa* reveals a regulatory effect of 3-oxo-C_12_-homoserine lactone in absence of its cognate receptor, LasR"

### SUPPLEMENTAL MATERIAL

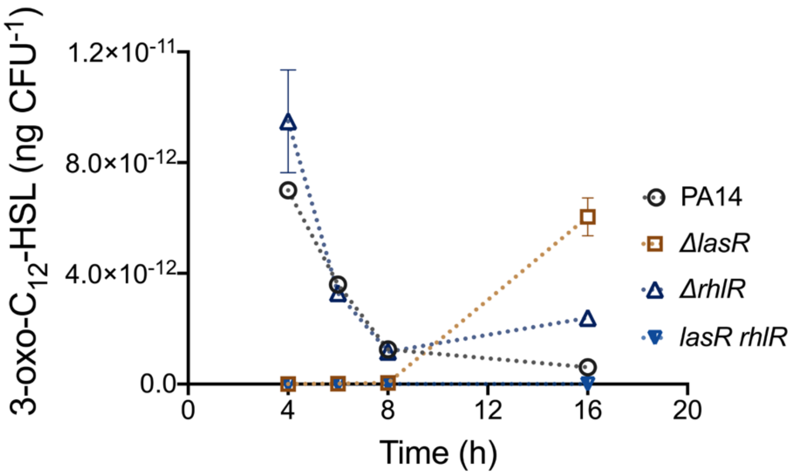


**Figure S1. Production of 3-oxo-$C_{12}$-HSL does require RhlR in LasR-active cells.** Concentration of 3-oxo-C12-HSL was measured at different time points during surface growth by LC/MS. Values were normalized by the viable cell counts and shown in ng $CFU^{-1}$. Values are means ± standard deviation (error bars) from three replicates.

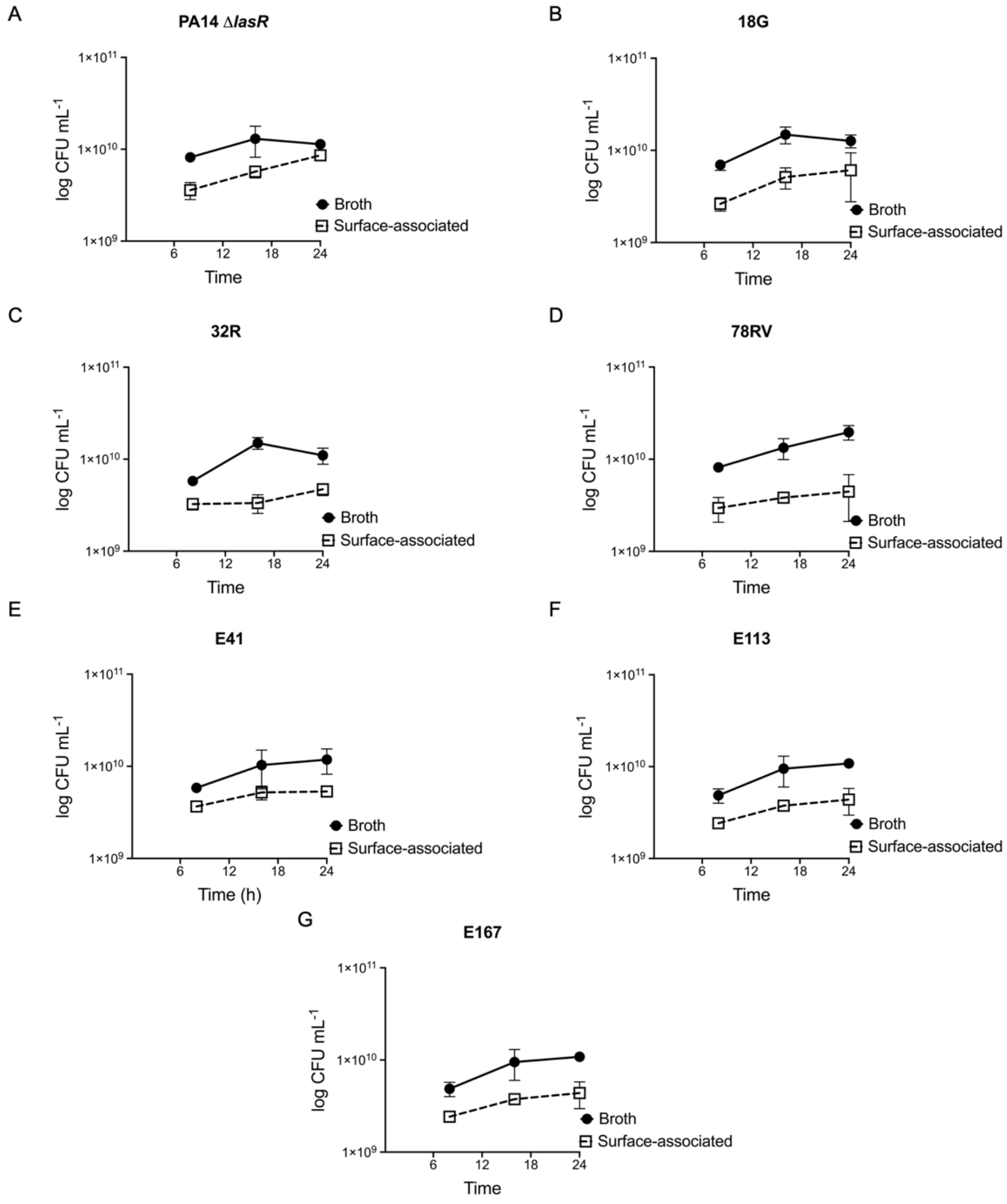


**Figure S2. Growth profile of natural occurring LasR-defective isolates.** Growth in broth and surface conditions was determined by the count of viable cells per millilitre (CFU $mL^{-1}$). This data is complementary to the one shown in Figure 4. The values are means ± standard deviation (error bars) from three replicates.

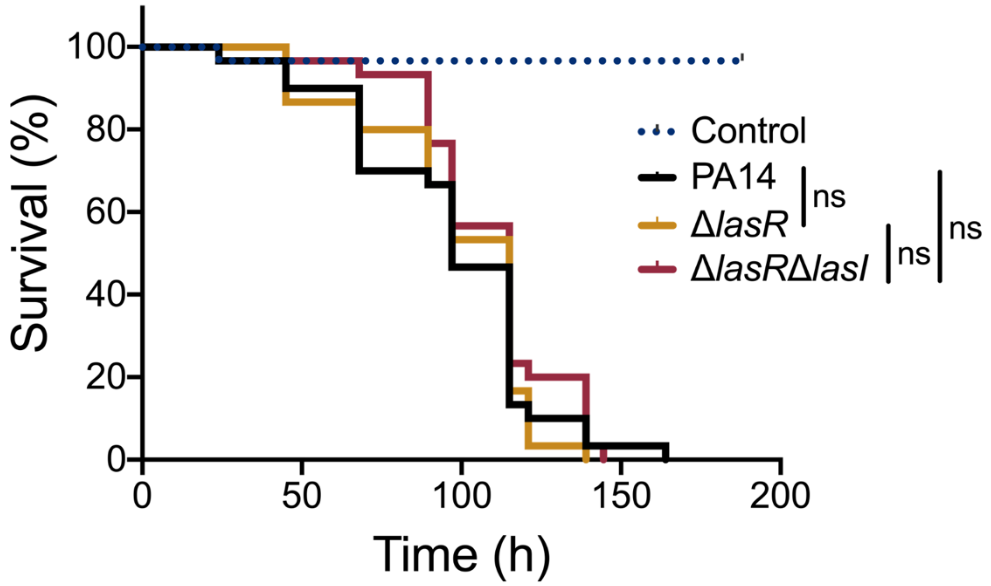


**Figure S3. Functionality of the *las* system is not required for *P. aeruginosa* virulence toward *D. melanogaster*.** Fruit flies were infected with suspended cells in 5% sucrose. Fly survival was monitored over time. *n* = 30 flies per group for each experiment. Experiment was performed independently twice. Statistical significance was determined using the Kaplan-Meier survival analysis. ns, non-significant.

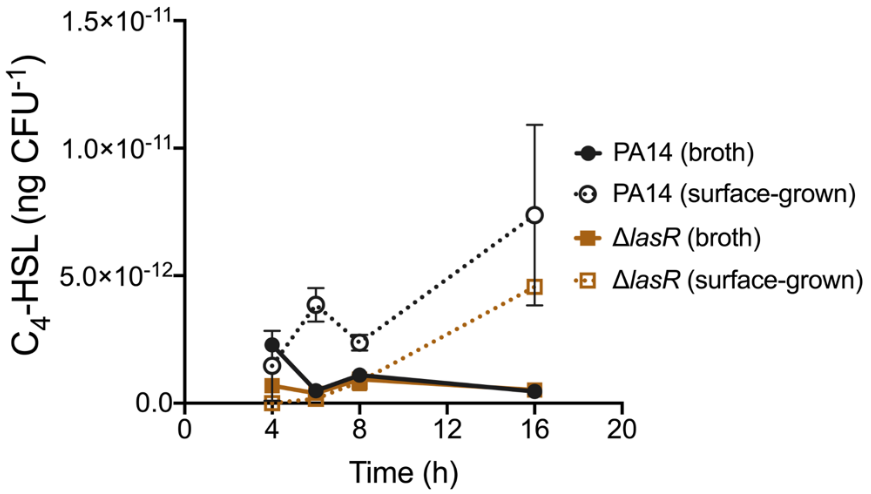


**Figure S4. Surface growth induces $C_4$-HSL production in both PA14 and its isogenic *lasR* mutant.** $C_4$-HSL concentration was measured in PA14 and the isogenic *lasR* mutant at different time points during planktonic (broth culture) and surface growth (surface of agar-solidified culture media) by LC/MS. Values were normalized by the viable cell counts and shown in ng $CFU^{-1}$. The values are means ± standard deviation (error bars) from three replicates.

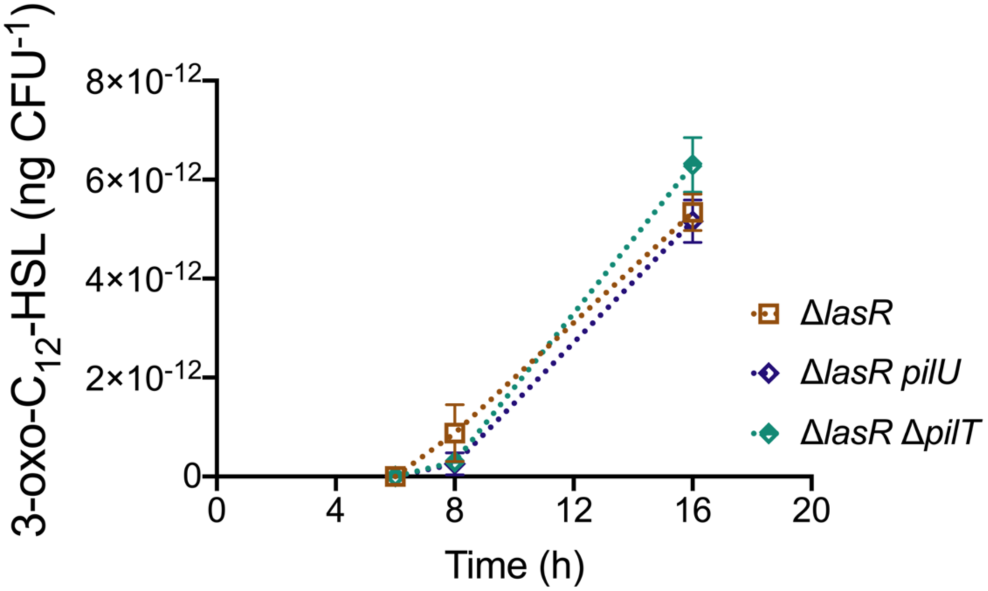


**Figure S5. Type IV pili motors PilU and PilT are not responsible for surface-primed 3-oxo-$C_{12}$-HSL induction.** 3-oxo-$C_{12}$-HSL concentration was measured in PA14 Δ*lasR* and the double mutants Δ*lasR pilU* and Δ*lasR* Δ*pilT* at different time points during surface growth by LC/MS. Values were normalized by the viable cell counts and shown in ng $CFU^{-1}$. The values are means ± standard deviation (error bars) from three replicates.

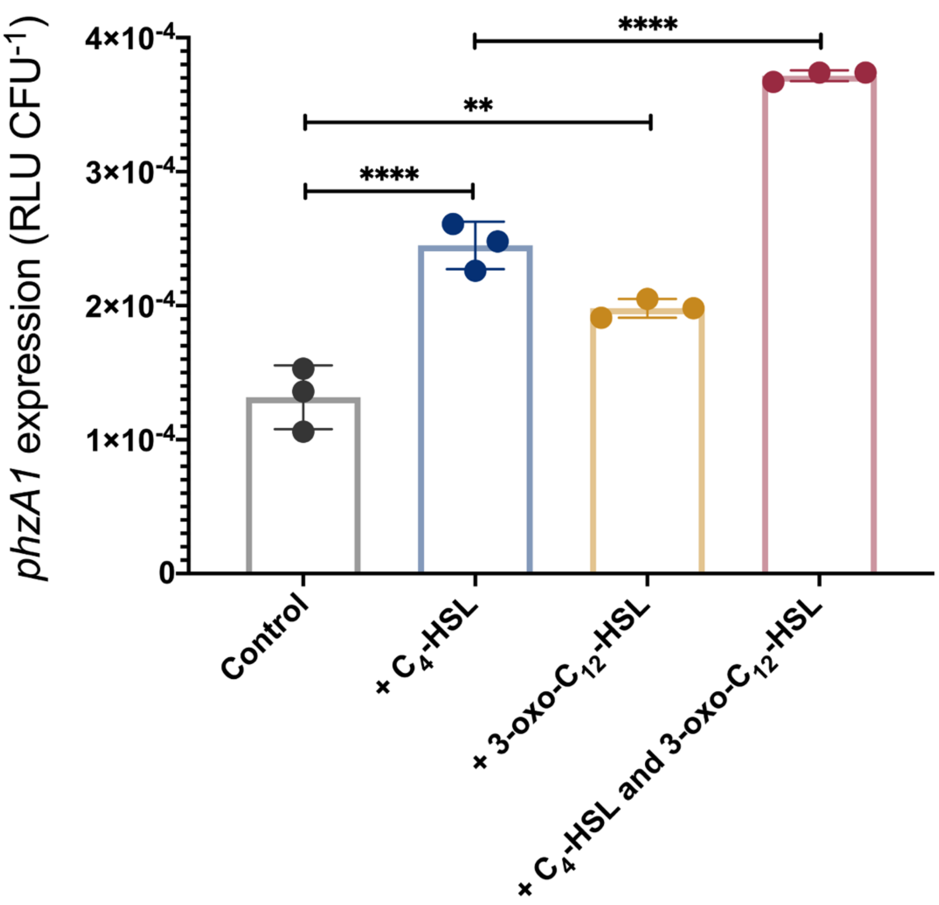


**Figure S6. Induction of the *phz1* operon by the 3-oxo-$C_{12}$-HSL is also seen with endogenous $C_4$-HSL.** Luminescence of the *phzA1-lux* chromosomal reporter was measured in a *las* system negative background (Δ*lasR*Δ*lasI*) after the addition of 1.5 μM of $C_4$-HSL, 3 μM of 3-oxo-$C_{12}$-HSL or both molecules at 8h. Solvent alone was used as control. Relative light units were normalized by by viable cell count and is shown in RLU $CFU^{-1}$. Statistical analyses were performed using one-way analysis of variance (ANOVA) and Tukey's multiple comparisons posttest with ** $P \leq 0.01$ and **** $P \leq 0.0001$.
